# Global Patterns and Drivers of Bee Diversity and Endemism on Islands

**DOI:** 10.64898/2026.02.28.708622

**Authors:** Leon Marshall, John S. Ascher, Robert J. Whittaker, Michael C. Orr, Alice C. Hughes, Julian Schrader, Patrick Weigelt, Holger Kreft, Nicolas J. Vereecken

## Abstract

Islands harbor a disproportionate share of global biodiversity^1^, yet insects, even invaluable pollinators such as bees^2^, remain underrepresented in island biogeography research^3^. Here, we present the first global checklist of island bees, recording 4,140 species across 306 islands. Although islands comprise only ∼5% of Earth’s land area, they support ∼20% of global bee diversity, and 43% of insular species are endemic, making up ∼8% of all known bee species.

Island bee species richness, mirroring continental trends^4^, peaks at mid-latitudes. Native richness increases with island area and declines with isolation, consistent with patterns in other taxa^5^. The strength of species–area relationships varies among biomes and is steepest in mediterranean-type systems, which also support disproportionately high bee richness relative to flowering plant diversity.

Endemism is highest on large tropical islands, reflecting extensive *in situ* diversification. Major centers of bee endemism include Madagascar, Malesia (e.g. the Greater Sundas and New Guinea), and Hawai’i, where a single large radiation of *Hylaeus* (*Nesoprosopis*) dominates^6^. Among islands capable of supporting endemic species, endemism scales strongly with total richness.

These findings highlight the need to integrate island bee diversity into global conservation planning and position bees as a model for understanding insect evolution and conservation on islands.

## Main Text

Islands are fundamental for studying patterns of biodiversity in time and space^1,7^. Their isolation and limited opportunities for colonization have fostered evolutionarily distinct and highly endemic flora and fauna^5,8,9^. Consequently, islands have long been central to fundamental questions in macroecology, evolution, community ecology, and species conservation^10^. Yet, despite their ecological and evolutionary importance, major gaps remain in our understanding of island colonization, evolution and diversification across animal taxa, especially arthropods.

Insects, the most diverse of all animal groups, remain underrepresented in global biogeographic studies, and conservation assessments^3^. This gap largely reflects logistical and taxonomic challenges of assembling standardized, high-resolution data for insects across geographically and politically fragmented islands that result in inadequate understanding of species’ distributions^11^. Consequently, tests of island biogeography theory have predominantly relied on vertebrates (especially birds), and land plants, while systematic studies across multiple archipelagos remain scarce both for insects and other invertebrates^12,13,14^.

Notable global efforts targeting other insect groups do exist, such as the Global Ant Biodiversity Informatics (GABI) database^15^, as well as efforts motivated by pest and pathogen spread, such as a global checklist of mosquito distributions^16^, and of patterns of island invasion by beetles^17^. Comparable progress has been made for Lepidoptera at regional scales, including recent large-scale analyses of butterfly biogeography across tropical Asian islands^18^ and New Guinea^19^. Nevertheless, comprehensive global syntheses, such as those available for plants^20^, of insect diversity and endemism across island systems are virtually absent.

There are approximately 21,000 described species of bees worldwide^21^. They play an important role in global ecosystems, pollinating the majority (87%) of wild flowering plants^22^, as well as many crop species^23^. On continents, bees exhibit a bimodal latitudinal diversity distribution, with peak richness in seasonally xeric mid-latitudes that support ecosystems with a mediterranean climate (hereafter referred to as mediterranean-type) ecosystems, as well as in adjacent desert and montane habitats^2,4,24^. This deviates from the latitudinal diversity gradient of most other groups of plants and animals which usually peak in the tropics^25^. While the continental-scale biogeographical and ecological patterns of bees are becoming better understood, it remains underexplored how they translate to island systems.

Generalizations about island bee biogeography have hitherto been rather qualitative in nature. For example, Michener^2^ noted that whereas very small bees might be carried over long distances by winds, wood-nesting bees may have dispersed across oceans in floating wooden rafts (timber). He suggested Polynesian transport by seafarers for certain wood-nesting bee taxa that reached some of the most remote oceanic archipelagos, such as Hawai’i, so bees detected by the even the first voyages of discovery cannot be assumed to be native.

Recent phylogenetic evidence indicates that several bee lineages colonized new regions during the Neogene (ca. 23–2.6 million years ago), *via* island-hopping, showing how stepping-stone dispersal may have shaped bee composition. For example, the families Apidae, Megachilidae, and Halictidae likely colonized Australia via movements from mainland Asia across the Indo-Malayan Archipelago^26^. Similar processes have been proposed to explain disjunct distributions in modern day Central America where ancient archipelago chains acted as stepping stones between North and South America before the continents joined^2^.

Conversely, disjunct distribution patterns in other bee groups appear to reflect their more limited capacity to disperse and subsequently colonize islands. For example, following the breakup of Gondwana, stingless bees (Apidae, tribe Meliponini) appear to have been unable to cross the substantial marine barriers that appeared, and their present Indo-Malay distribution likely reflects episodes of sea level fluctuation and transient land-bridges, rather than long-distance overwater dispersal^27^.

These examples indicate the importance of accurately documenting insular bee diversity to achieve a more complete and fine-grained understanding of global bee biogeography and conservation. Yet, despite this need, comprehensive data remains sparse. Only 542 bee species have been assessed on the global IUCN Red List, including just 240 species recorded from islands and 68 island endemics, of which 26 are considered threatened, and for many of these, distributional data are not publicly accessible^28^. The limited representation of island taxa on global conservation lists does not reflect the well-established vulnerability of island biotas: for example, the majority of U.S. federally listed endangered bee species are native Hawai‘ian endemics^29^.

Here, we present the first global assessment of native bee diversity on islands. Specifically, we investigate: (i) global patterns of native bee richness and endemism; (ii) the influence of island area and isolation on native bee richness; (iii) the drivers of endemism and how endemism scales with richness; and (iv) the relationship between insular bee diversity and angiosperm richness. We expect the patterns of insular bee richness and endemism to be shaped by area and isolation, as predicted by island biogeography theory^7^, but also by biotic and climate constraints and associations, including mutualistic relationships with flowering plants^22^.

By addressing these questions, our analyses fill knowledge gaps, providing the first global baseline for investigating the evolutionary and biogeographic history, ecology, vulnerability, and ecological roles of island bees, while highlighting priority regions for future research and conservation at both archipelago and single-island scales. More broadly, this baseline enables rigorous tests of major hypotheses in island biogeography and (meta-)community assembly^1^ using a taxonomically diverse, and ecologically and economically important group.

### Global distribution of bees on islands

Our final dataset consisted of 11,171 unique island−native species combinations, representing 4,140 native species across 306 islands (See Methods for details of this classification) (Fig. 1a). Consequently, of the approximately 21,000 bee species described worldwide, our data suggested that 20% occur on islands. Of these, 43% (1,725 species) are restricted exclusively to islands, accounting for ∼8% of all described bee species. Although this proportion is lower than that reported for vascular plants, for which 21% of the global flora are island endemic^9^, it nevertheless underscores the disproportionate contribution of islands to global bee diversity relative to their area, approximately 5% of global land mass^1^. Although we focus here on native bee richness, we also noted introduced species. We identified at least 43 species as introduced on 142 islands, with *Apis mellifera* being the most widespread, introduced to 122 islands in our dataset (see Table S1). Invasion due to anthropogenic drivers is a major threat to insular diversity^30^, and evidence, at least for *A. mellifera*, suggests negative effects of invasive bees on native biodiversity on islands (e.g.^31^).

**Figure 1.**
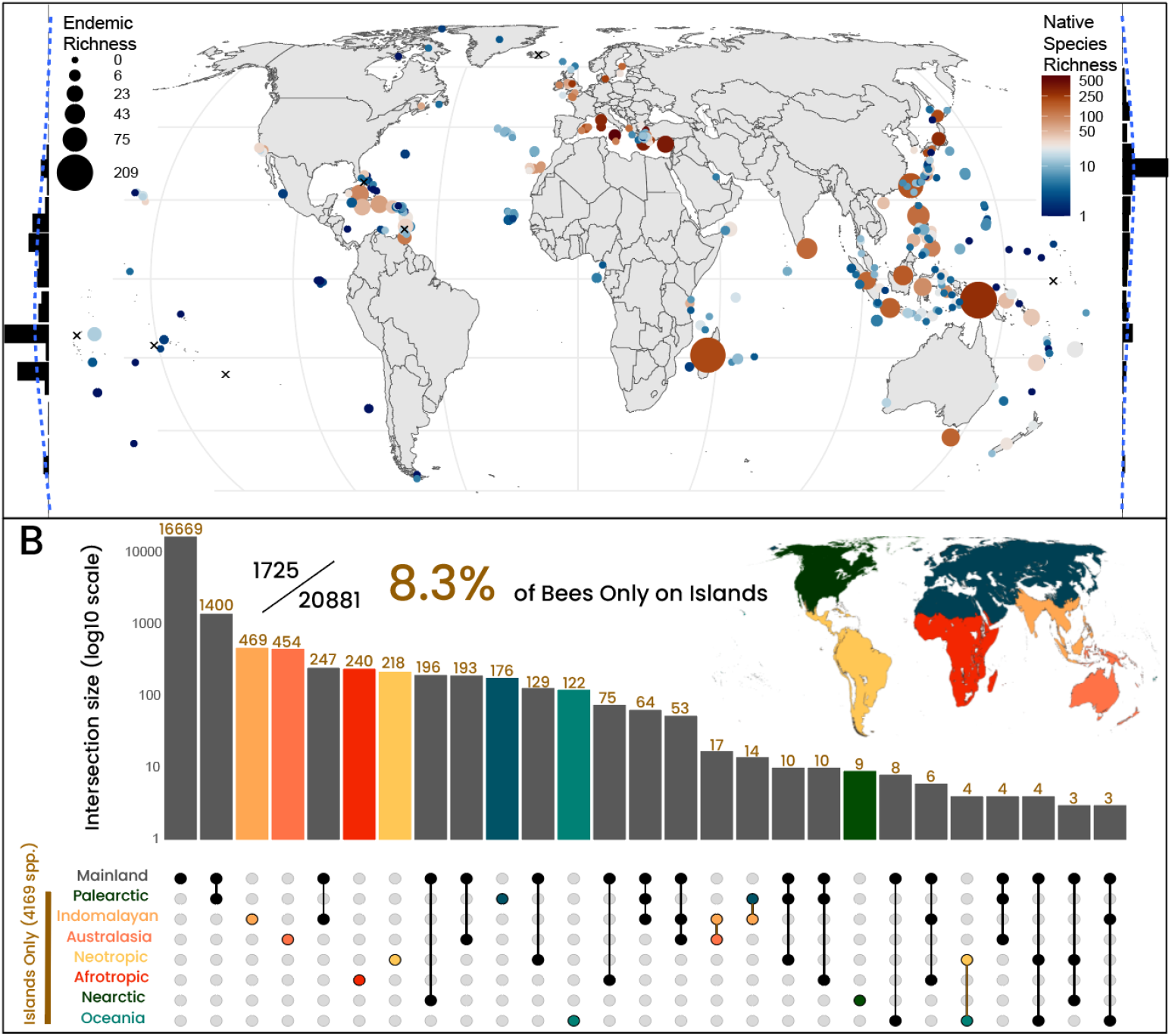
Global distribution of bees on islands. (A) Map of global bee diversity on islands. Each circle represents an island for which there are recorded native bee species. Color gradient is log-scaled and indicates the total number of native bee species recorded for each island; the size of the point or circle is proportional to the total number of endemic species found on each island. Endemism is measured as the total number of island (or island group, see Methods) endemic species. The histogram to the left of the figure shows the total number of endemic bee species by latitude and the histogram to the right shows the total number of bees on islands by latitude (5 degrees latitude steps; native richness max = 3,562 species (35 — 40°) and endemism max = 294 species (-10 — -5°). Note this pattern does not control for variation in the number and size distribution of islands across latitude, but see below (Fig. 2). Islands marked with an X have recorded bee species consisting exclusively of non-native species. (B) Upset plot showing global distribution of bee species endemic to and shared between islands in different zoogeographical realms^33^ and the Mainland. Of 4,169 bee species found on islands, 1,725 are recorded only from islands, representing a 8.3% share of global bee diversity unique to islands. Each bar shows the number of species unique to that geographic combination. Colored bars and colored realm names represent the number of species endemic to islands within that realm. Colored circles indicate island-only comparisons. Gray bars indicate species shared across the island realms or mainland.

Bee species richness peaked in the Western Palearctic Realm, whereas endemism (species only found on islands) was highest on Indomalayan (501 spp.) and Australasian (472 spp.) islands (Fig. 1b). Clear differences in island bee richness were also apparent among biomes (Fig. S1), however this pattern was influenced by variation in island area. Below we show how these patterns changed across area and biome. Species richness was highest in mediterranean-type biomes: Sicily (636 spp., ∼25,000 km^2^), was identified as the most species-rich island worldwide, and Cyprus (384 spp., ∼9,300 km^2^), the most species-rich oceanic island, as well as the East Aegean islands (592 spp., ∼5,000km^2^) supported large bee faunas dominated by widespread continental species, reflecting their limited degree of isolation. In contrast to the high richness, endemism on these islands was low, with only Crete (10 spp.) and Cyprus (23 spp.) showing visible levels of island endemism (Figure S1). Despite its proximity to the Levant and its complex origins in tectonic uplift, Cyprus was treated here as an oceanic island due to its lack of historical land-bridge connections to the mainland. The most species rich oceanic island of strictly volcanic origin is Tenerife (77 spp., ∼2,000 km^2^).

Tropical islands, by contrast, harbored lower overall native species richness compared to mediterranean-type islands but markedly higher levels of endemism. New Guinea supports 286 species of which 209 (73%) were endemic, making it by far the richest island in the Australasian realm. The nearby Maluku Islands archipelago hosts 102 species, including 62 endemics (61%, ∼34,000 km^2^), and contains Seram, the most species-rich tropical oceanic island for bees (87 spp., ∼18,000 km^2^).

In the Neotropics, Hispaniola stood out as the most species-rich oceanic island, with 68 species. In the Nearctic, the predominantly oceanic California Channel Islands had a high overall richness (156 spp., ∼910 km^2^), but low levels of endemism (5%). In the Afrotropics, Madagascar stood out with 212 species, of which 187 (88%) were endemic. Within Oceania, which consists only of oceanic islands, the Hawaiian Islands emerged as a major hotspot, with 71 native species of wild bees, 64 (90%, ∼17,000 km^2^) of which are endemic to the archipelago, all belonging to a single lineage of *Hylaeus* (*Nesoprosopis*)^6^. Among other oceanic systems, the mediterranean climate Canary Islands hosted a similarly high number of endemic species (52 of 139 species), though at a lower proportional level of archipelago endemism (37%, ∼5,400 km^2^).

Globally, bee richness on islands peaked in the Northern hemisphere at mid-latitudes (Fig. 1a), mirroring the mid-latitude peak of richness observed on mainlands^4^, although this pattern was not controlled for variation in the number and size distribution of islands across latitude, but see below. Furthermore, the limited number of mediterranean-type islands in the Southern Hemisphere in our dataset meant that we did not observe an island-only bimodal distribution pattern comparable to that observed on mainlands^2,4^. This pattern occurred both on continental and oceanic islands, although many of the larger islands within the mediterranean-type biome were continental or complex in origin. Such environments favor diverse wild bee assemblages through a combination of evolutionary history and ecological conditions. Bees are thought to have originated in the arid environments of Western Gondwana^2,26^, with limited but predictable rainfall that generates short yet intense blooming periods and a high flowering plant diversity in open habitats^2,4,32^. These conditions promote temporal synchrony between bee flight periods and floral resources, fostering increased specialization and resource partitioning that reduces competition^2.^ Our global results support these conclusions but trait-based research focusing on the biome-specific patterns of habitat and dietary specialization of bees on islands are needed to directly test these claims.

## What drives the observed patterns of richness?

To test fundamental aspects of MacArthur and Wilson (1967)’s theory of island biogeography for bee diversity, we employed a mixed-effects model with archipelago identity as the random effect. The final best model was selected based on Akaike Information Criterion corrected for sample sizes (AIC_c_) and included island area (log_10_km^2^) and its interaction with biome, as well as isolation (log_10_ distance to continental mainland in km) as fixed effects. This model explained 65% of the variance in species richness through fixed effects (area, isolation, island type and biome), and 75% when including archipelago identity (conditional R^2^). This moderate difference in R^2^ values indicates that native richness is strongly governed by broad biogeographic variables, with some unexplained regional structure.

Island area showed a consistent positive effect on native bee richness but the island species-area relationship (ISAR) slopes differed significantly among biomes (Fig. 2a and Table S2). For every 10-fold increase in island area, mediterranean-type biomes showed the strongest species–area response, with bee species richness increasing by approximately 90% (*β* = 0.50), this was significantly steeper than all other biomes (Table S3). Tropical islands exhibited an intermediate effect, supporting about 40– 45% more species per tenfold area increase (*β* = 0.28), also steeper than all other biomes, except mediterranean-type. Temperate islands showed a more modest increase of approximately 25–35% (*β* = 0.19). In contrast, desert islands displayed only a ∼10% increase in richness per tenfold increase in area (*β* = 0.01), indicating that island size is not a major constraint on bee diversity in these biomes. It may be noted that Socotra forms something of an outlier in that it supports a comparatively high richness (26 native species) and high endemism (∼60%) compared to other desert-biome islands, likely due to its geographic positioning with bee fauna origins from both the Palearctic and Africa^34^. Boreal islands similarly exhibited a very weak area effect (*β* = 0.05). The impact of biome on species area slopes was observed more clearly on oceanic islands (Fig. S2 and Table S3), specifically, the species-area slope for Mediterranean-type islands was significantly greater than the others. This was not the case for continental islands where these species-area slope differences were far less pronounced (Fig. S3 and Table S4).

**Figure 2.**
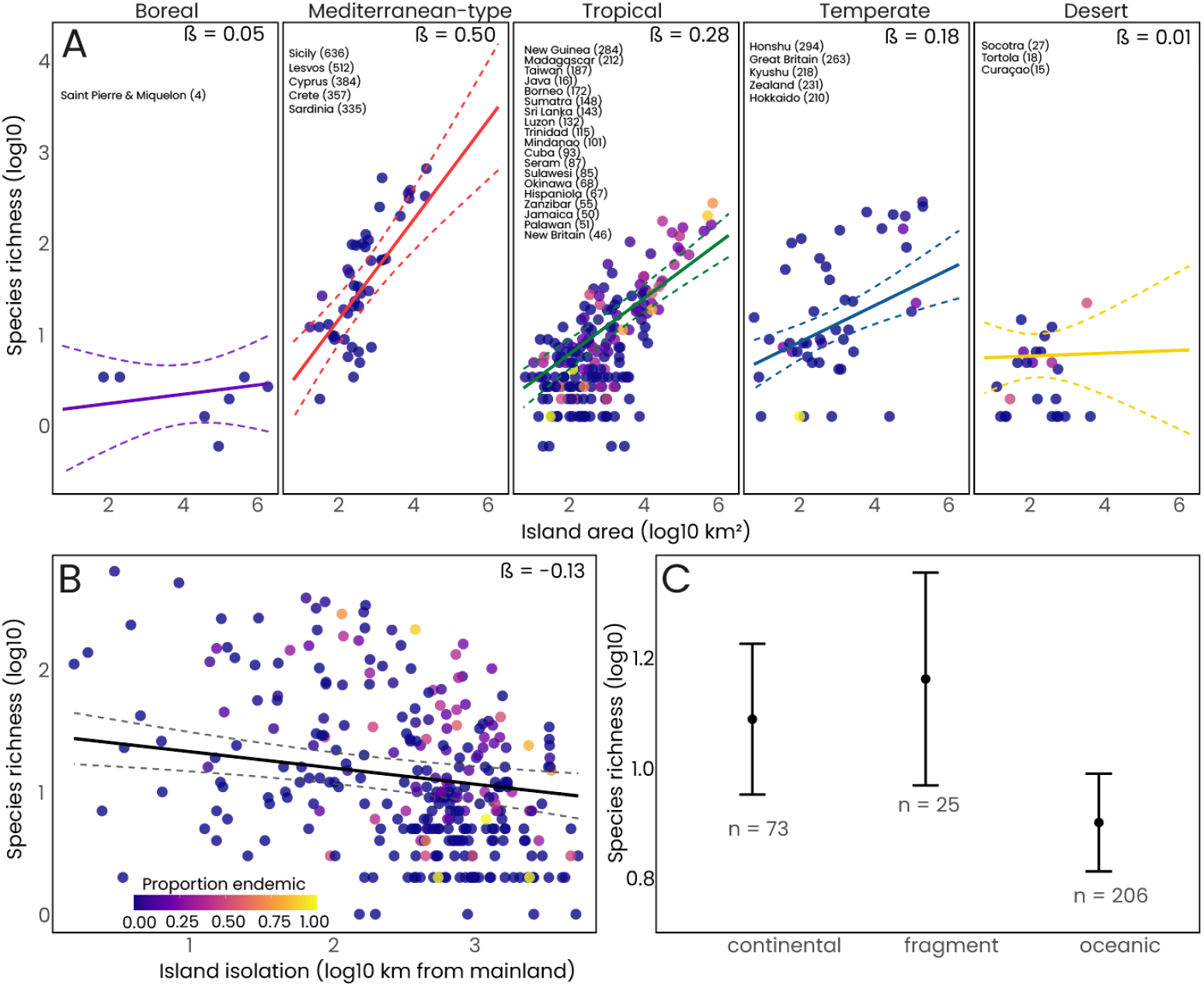
Drivers of island bee species richness. (A) Biome-specific species–area relationships. Solid lines represent fitted slopes from mixed-effects models, with dashed lines indicating 95% confidence intervals. Slopes (β) are reported for each biome. Point colour indicates the proportion of endemic species per island. Labelled islands are the top 10% most species-rich islands within each biome (values in parentheses show total native bee richness). (B) Relationship between island isolation (log_10_ distance to mainland) and native bee species richness. Point colour indicates the proportion of single island endemics (SIE). (C) Predicted bee species richness by island type (continental, fragment, oceanic) from the full mixed-effects model, holding other predictors constant (predictions represent an island of median area (∼440 km^2^), median isolation (∼610 km from mainland), and median sampling coverage (∼36% after sample-size adjustment, raw median 79%). Circles indicate the model-predicted means ± 95% confidence intervals, with the number of islands shown below each category.

Richness decreased with island isolation (β = -0.13, *p* < 0.001) (Fig. 2b and Table S2), corresponding to a reduction of roughly 15% in species richness for each tenfold increase in distance (km) from the mainland. This isolation effect was uniform across all biomes, supporting the fundamental principle that remote islands receive fewer colonists and as a consequence maintain lower richness as a function of their area^7^. We also observed a clear influence of island type on richness. Oceanic islands supported substantially fewer species than continental or fragment islands (β = -0.19, *p* = 0.006), consistent with their more limited historical species pools and generally younger geological ages^1^.

Our results supported the conclusion that ISARs can vary considerably as a function of island geo-environmental properties^12,35^. Specifically, bees may benefit from biome-specific increases in ecological complexity in mediterranean-type biomes due to temporal differences in habitat availability, which can permit increased richness even at small scales, resulting in area having a stronger effect on richness than in tropical or temperate biomes^2,4^. The finding that mediterranean-type islands in our models have the steepest ISAR slopes and highest richness values, align with the high seasonality of the climate and flowering plant diversity characteristic of mediterranean climates^36^, providing diverse nesting sites and floral resources across temporal scales.

While biomes explain considerable variation in richness on islands, they remain broad proxies for climate and habitat diversity. The steep ISAR slope for mediterranean-type systems suggests that capturing resource availability and diverse microhabitats may yield finer-scale insights into the mechanisms underlying island bee biogeography. Habitat diversity and environmental heterogeneity, and the potentially associated diversity of mutualist partners, constitute important predictors of species richness^37^, which appears to be reflected in our model by both area and biome. For many large islands with a high elevational range, like Madagascar, Tenerife, New Guinea, or Hawai’i’ (big island), representing the island as a single biome likely underestimates its pronounced environmental heterogeneity, which has presumably contributed to both high species diversity and exceptional levels of endemism. Additionally, regional geographic context is an important consideration, as shown by the islands of Lesbos and Cyprus, which show comparatively elevated species richness values for their area (positive residuals from the mediterranean-type ISAR; top 5% among mediterranean-type islands). These islands sit at the border between the European bee fauna and the exceptionally rich western Asian fauna, and the resulting high comparative species richness for their area may be due to this dual species pool from which bees can colonize^38^.

Sampling effort was uneven across regions, as expected for a global dataset. Substantial variation in sampling completeness existed across the dataset (Fig. S4), with parts of Oceania, Indomalaya and Australasia being comparatively under-sampled. To account for this heterogeneity we included coverage (the proportion of individuals belonging to observed species and adjusted for sample size) as a fixed effect in the native richness model (Table S2). Coverage had a positive effect on richness (β = 0.22, *p* < 0.001). Islands with a greater sampling effort tended to have higher estimated richness. However, controlling for effort did not weaken the central results above. The island-biome-area effects remained strong after adjusting for sampling completeness (Table S5), indicating that the relationships between area, isolation and richness were not a result of uneven sampling effort. Future efforts to digitize and provide open-access to museum collections^39,40^, conduct targeted expeditions to historically difficult-to-sample islands^2,41^, and harness citizen science platforms such as iNaturalist^42^, will be critical to resolving knowledge gaps in island bee diversity.

### What drives bee endemism on islands and does it scale with richness?

To identify the ecological drivers of endemism and assess whether richness can explain endemism patterns, we fit the same linear mixed-effects models as for native richness (Fig. 3a b). Across islands capable of supporting single island endemics (SIE>0, n= 101), area was the only consistent predictor of SIE richness (β = 0.35, *p* < 0.001, R^2^ = 0.50; Table S6). Unlike patterns of native richness, where area effects varied significantly among biomes, SIE showed a uniform ISAR across all biomes. Isolation had only a weak, non-significant negative influence (β = -0.06, *p* = 0.36; Fig. 3a b) and sampling effort appeared in best models but had no significant effect on SIE (Table S7), though we note that it could bias endemism estimates in both directions, oversampling endemism on well-studied islands where the same species have not yet been recorded elsewhere, and under-sampling where undiscovered endemics likely remain to be sampled.

**Figure 3.**
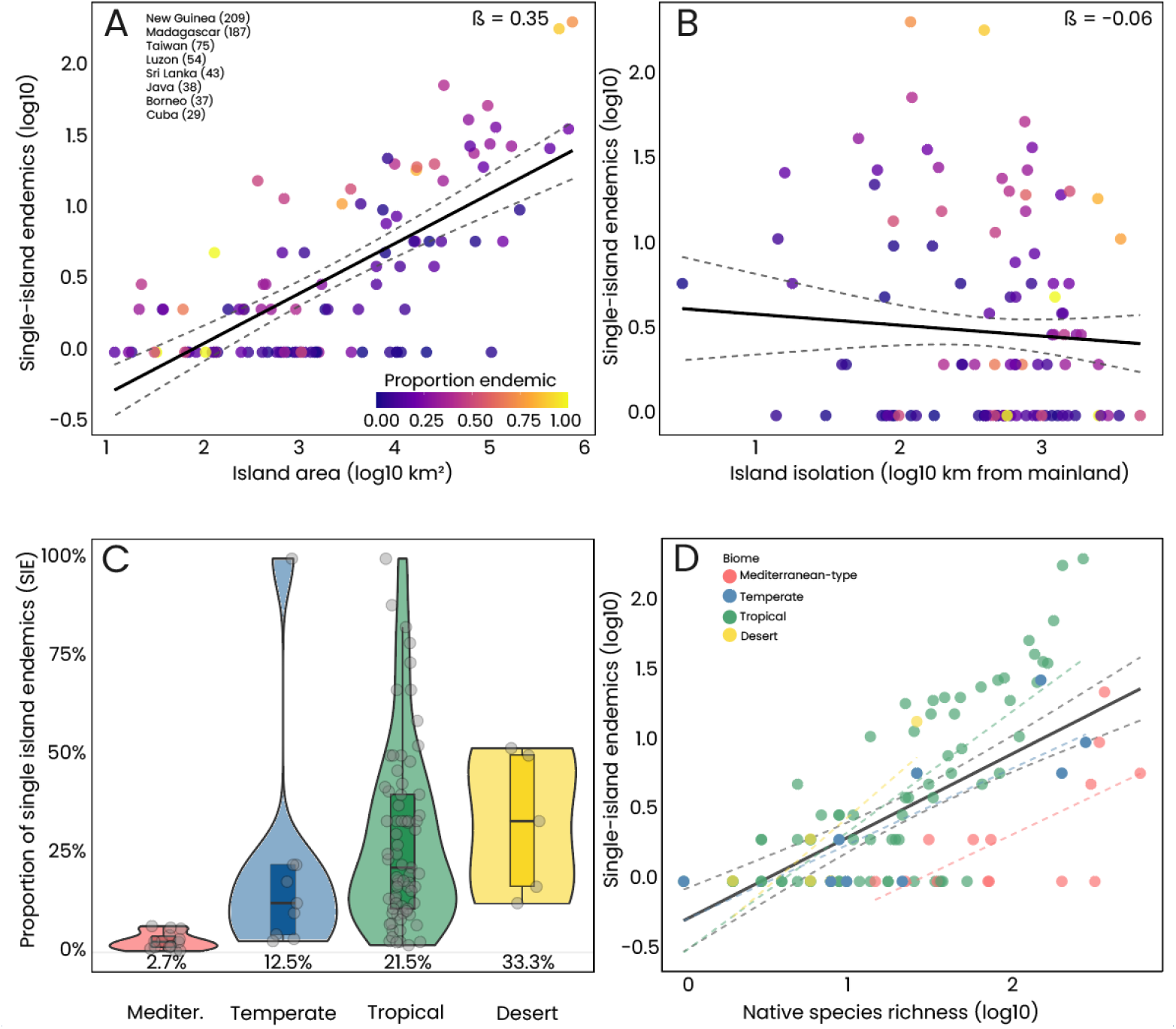
Drivers of island bee endemism and spatial relationship with richness patterns. (A) Relationship between island area (log_10_ km^2^) and single island endemic (SIE) species richness for islands with at least one single-island endemic (n = 101). Black line shows the overall trend with 95% confidence interval (gray shading). Circles are colored by the proportion of native species that are endemic. Islands with highest SIE richness are labeled with the number of SIEs in parentheses. (B) Relationship between island isolation (log_10_ distance to nearest mainland, km^2^) and SIE richness. The Black line shows the overall trend with 95% confidence interval. Circle colors as in panel A. (C) Distribution of endemism rates (proportion of native species that are endemic) across biomes. Violins show full distribution; horizontal lines mark 25th percentile, median, and 75th percentile. Individual islands are shown as circles. Biomes ordered left to right by median endemism rate. Median values displayed below each distribution.(D) Relationship between total native species richness (log_10_) and SIEs richness (log_10_), colored by biome. Dashed black line shows overall linear relationship; dashed colored lines show biome-specific relationships. All panels include islands with at least one single-island endemic (SIE ≥ 1; n = 101).

The observed patterns were consistent across data subsets. On oceanic islands with endemics (n= 59), area remained the strongest predictor (β = 0.25, *p* <0.001, R^2^ = 0.28; Fig. S5, Table S6). Among islands high in SIE (>5% of native richness; n= 81), SIE richness increased with area (β = 0.39, *p* < 0.001, R^2^ = 0.65) and decreased marginally with isolation (β = -0.15, *p* = 0.05) (Fig. S5; Table S6). These results paralleled global patterns of vascular plant endemism on islands^43^. The mid-latitudinal peak in bee species richness^2^, seen on islands predominantly in the Northern Hemisphere, did not translate into increased bee endemism. Rather, bee endemism was greatest in the tropics (Fig. 3c). Model fit of models predicting SIE from overall native richness alone improved significantly with the inclusion of biome (ΔR^2^ = 0.18, χ^2^ = 33.1, df = 3, p < 0.001; Table S8). Mediterranean-type islands supported significantly fewer SIEs than all other biomes (Fig. 3d; Fig. S6) indicating a partial decoupling between richness and endemism. This contrasts with vascular plants, for which the richest islands also tend to host the highest numbers of endemic species, particularly large tropical islands^9^. Our results were consistent with earlier suggestions that the relationship between richness and endemism on islands requires further exploration and varies among taxa^44^.

Patterns for archipelago endemics (AE) were broadly similar. Across islands hosting AE (n = 215) area was again the main predictor (β = 0.22, p = 0.03). However, unlike SIE, AE richness included the interaction between area and biome, with tropical islands showing the steepest area-AE slopes (β = 0.38) and desert islands the shallowest (β = 0.05; pairwise comparison: p = 0.01; Table S9). This shallow slope for desert islands was driven almost exclusively by the Galápagos, which are comparatively large desert islands that host a single archipelago endemic species across the archipelago (*Xylocopa darwini*). As with SIE, including biome improved models predicting AE from richness alone (ΔR^2^ = 0.12, χ^2^ = 40.04, df = 3, p < 0.001; Table S8). Mediterranean-type islands again supported fewer AE than expected (Fig. S6), although the overall difference was less pronounced; for example the Canary Islands showed AE richness consistent with their overall richness (Fig. S6). This likely reflects that AE captures more ancient diversification on islands compared to SIE^45,46^.

Together, these patterns reveal that bee endemism is concentrated on large tropical islands and certain oceanic archipelagos rather than in areas of peak richness globally. New Guinea and Madagascar stand out for their high raw numbers of endemic species whereas high proportions of endemic species characterize island systems including the Caribbean, the Philippines, and Hawai’i. These patterns reflect include distinct evolutionary histories, high endemism in Madagascar likely reflects multiple independent radiations whereas in other systems single radiations drive high proportions of endemism, such as *Hylaeus* (subgenus: *Nesoprosopis*) in the Hawai’ian archipelago^6^ and *Lasioglossum* (*Homalictus*) across Pacific islands such as Samoa and Fiji^47,48^. Conversely, the factors that support high richness in mediterranean-type islands, nearby continental source pools and high habitat diversity across seasonal gradients likely inhibit the isolation needed for *in situ* diversification resulting in lower endemism^49^. At a global scale, bees reinforce the idea that endemism, and by extension conservation threat, is not tightly linked to overall species richness or richness-based biodiversity hotspots, as some island regions support disproportionately high levels of bee endemism despite comparatively low species richness^50^.

### Does bee diversity track global angiosperm diversity?

Building on these findings, we additionally examined how plant diversity, as a potential biotic driver, covaries with bee richness, given that bees depend strongly on flowering plants for food and nesting resources^22,24^. Across the 181 islands for which we had both angiosperm richness data and native bee richness, we observed a positive correlation (β = 0.95, *p* < 0.001; *R*^*2*^ = 0.50; Fig. 4a and Table S10). However, the relationship varied significantly across biomes (Kruskal–Wallis *χ*^*2*^ = 59.98, *p* <0.001; Fig. 4b, Table S10). Islands within mediterranean-type and temperate biomes showed consistent positive deviations, harboring 5.1x and 2x more bee species than expected based on angiosperm diversity alone. In contrast, islands with tropical biomes had fewer (0.7x) bee species than predicted by the model.

**Figure 4.**
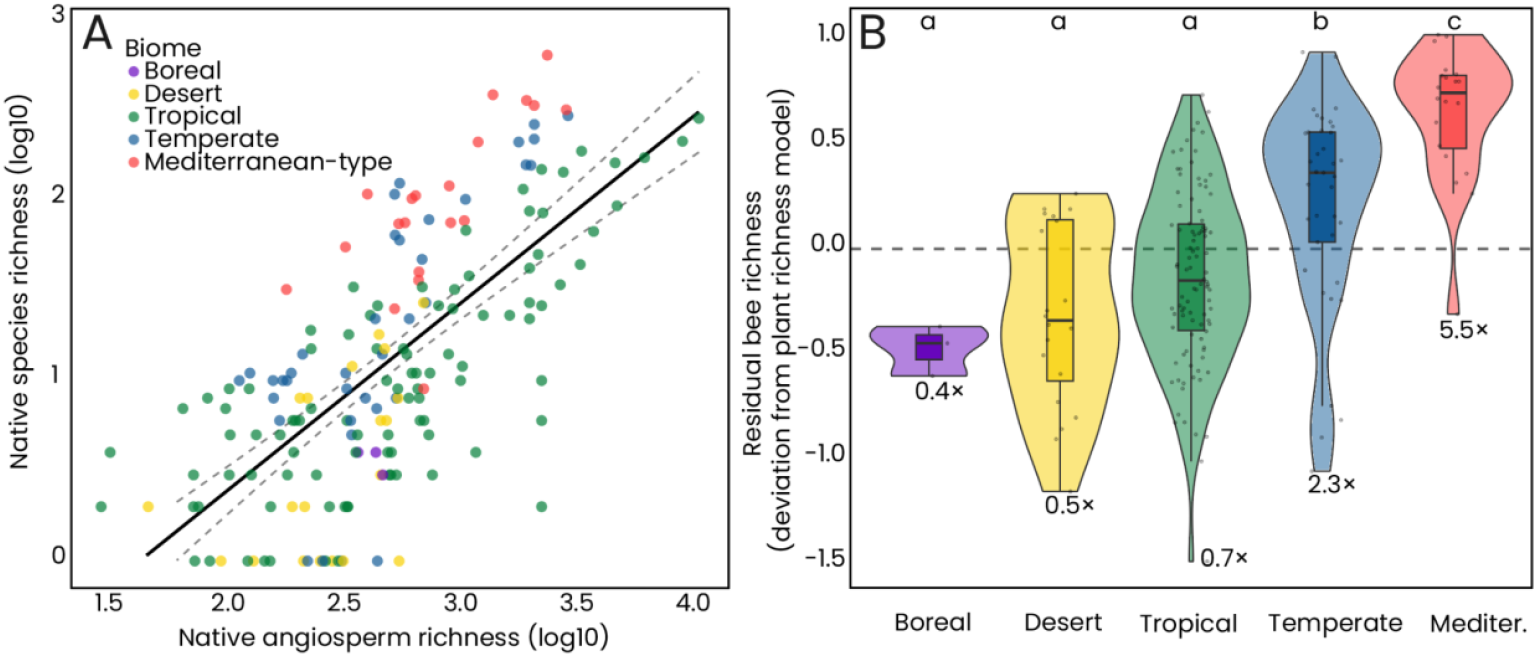
The relationship between native bee richness and angiosperm richness across islands and the differences between biomes. (A) The relationship between native angiosperm richness (log_10_) and bee species richness (log_10_) across islands worldwide. The solid black line represents the fitted linear model, with shaded bands indicating 95% confidence intervals. Circles are coloured by biome. (B) Residuals from the bee–plant richness model, representing deviations in bee richness relative to that expected from plant richness alone. Residual distributions differ significantly among biomes (Kruskal–Wallis test; Table S10). Letters indicate whether biomes differ significantly, biomes which share a letter do not differ significantly (p ≥ 0.05).

The relatively weak alignment between plant and bee richness across biomes further supports the theory that temperature seasonality, not primary productivity or absolute floral richness, is the dominant driver of bee specialization, diversification and ultimately species accumulation^51^, in line with the results for richness and endemism. Seasonal climates promote bee specialization through temporal resource partitioning while perhumid tropical systems, despite high plant richness, present ecological filters that limit bee diversity. These results also suggest that mutual ecological dependence does not necessarily produce parallel biogeographic patterns when distinct abiotic constraints shape the broader evolution of the groups. Therefore, capturing the complex networks and communities that wild bees form with flowering plants is crucial for a deeper understanding of ecological drivers on islands^52^. For example, recent evidence shows that flowering plant communities can still rely on bee pollinators even when bee richness is comparatively low, further supporting that floral diversity alone does not dictate bee diversity across biomes^53^.

### Conclusions and future research

Our study demonstrates that islands harbor a substantial share of global bee diversity, including many species found nowhere else. We found that species richness and endemism are partly decoupled, highlighting that these two metrics capture distinct aspects of insular bee diversity. Richness peaked in mid-latitude, mediterranean-type islands in the Northern Hemisphere whilst endemism peaked in large tropical islands, with these differences being strongest for oceanic islands. Our findings reveal that island bee endemism is largely decoupled from richness-based hotspots, particularly in tropical biomes where extinction risks are highest but formal assessments are most lacking. This spatial discordance underscores an urgent need to shift global conservation priorities toward these ‘evolutionary reservoirs’ to prevent silent co-extinction cascades within unique insular pollinator networks. These patterns were further supported by the biome-dependent coupling between angiosperm and bee richness, showing that floral richness alone does not predict bee diversity across islands.

Despite the advances presented here, much remains to be explored if we are to understand the mechanisms behind the patterns of colonization and speciation, the potential for new discoveries and the threats facing these endemic island bees. An increasing number of molecular phylogenies opens up new research avenues for addressing the evolutionary history of bees^26,54^. Combined with increasingly detailed knowledge on distribution patterns and traits^55^, this provides the potential to explore patterns of historical colonization, extinction and speciation on islands. While we provide clear evidence that area and isolation are important drivers of bee diversity and endemism, we also highlight that the richness patterns mirror, at least in the Northern hemisphere, the global distribution pattern of wild bees on continents^4^. Further models could build on our results here by including additional predictors related to ontogeny, habitat diversity, fine-scale climate data, and anthropogenic pressures.

Bees are particularly well suited as a model taxon for island biogeography. They are (i) pivotal pollinators, (ii) relatively straightforward to survey using standardized field methods, and (iii) they are closely linked to plant reproduction and ecosystem functioning^22^. Expanding this study and our dataset and integrating phylogenetic, ecological and genomic approaches will help clarify the evolutionary history of island bees and guide conservation efforts of these essential, yet often neglected pollinators.

## Methods

### Global checklist of bees on islands

We sourced the presence of bees on islands based on the information available on Discover Life and more detailed and recently updated information, notably for type locality and collection event data, extracted from source files for that resource^21^. These expert-vetted lists were filtered for island records and joined with spatial data to align species presence with island identifiers. The checklist data on DiscoverLife were collected from a variety of sources, including taxonomic revisions, primary species descriptions (including synonyms), verified human observations, and undigitized museum records. To avoid complications from historical boundary changes, species were recorded at the current state or country level. The data are available online as a matrix at the country level, with species-specific details provided for finer geographic scales. For further details on how the checklists were sourced see^4^. For islands not clearly associated with a specific country or state/province, additional sources were used to confirm species presence based on the broader political region. We then cross-referenced specimens with island-specific faunal checklists from the published literature; for example, when a species was known only from an archipelago we tried to determine its presence on specific islands within the archipelago. These included studies from Abd al Kuri and Socotra from the Socotra Archipelago^33^, the Azores including Corvo, Faial, Flores, Graciosa, Pico, Santa Maria, São Jorge, São Miguel, and Terceira^56^, Balearic Islands including Ibiza, Mallorca, and Menorca^57^, Bonaire^58^, Cape Verde including Boa Vista, Brava, Sal, Santiago, Santo Antão, São Nicolau, and São Vicente^59^, Cuba and East Grand Bahama^60^, Fiji^46,61^, Isla de Juventud^62^, Juan Fernández Islands^63^, Madeira Archipelago including Madeira, Porto Santo, and Deserto Grande, and Selvagens Islands including Selvagem Grande^64^. European island checklists were extracted from the latest checklists for European species from Reverte et al.^65^ and included Åland islands, Great Britain, Ireland, Iceland, Faroe Islands, Malta, Corsica, Sardinia, Sicily, Crete, Cyprus, and Isle of Man. We also included validated iNaturalist records (up to December 2024) for all islands available from the initial checklists. The checklists for the Greek islands of Corfu and Rhodes were expanded with records from Rhodes^66,67^, Corfu^68^ and the personal collection of TJ Wood. Recent additions to Corsica and Sardinia were also added^69^. Names were cleaned using the synonym lists from Orr et al.^4^ and Dorey et al^70^. In some cases, it was not possible to separate archipelago level data into island records and some archipelagos were therefore kept in the dataset (e.g., for the Solomon Islands).

To classify non-native species and calculate final island native species richness, we used Russo^71^ as a baseline, complemented by existing checklist information and expert opinion from co-author J.S.A. Using the check-lists we calculated three measures of wild bee richness and endemism on islands. They included: (i) native species richness, calculated as the total number of unique native species included in the checklist for each island (or archipelago, as the case may be); (ii) single island endemism, calculated as the total number of species on the checklist only known from that island; (iii) archipelago endemism, calculated as the total number of species found only on that archipelago: this includes all single island endemics.

### Spatial information on islands and island selection

Using the above checklists we extracted a spatially explicit database of islands by matching the names from the checklist to those found in the global islands (greater than 1 km^2^ in area) database of Sayre et al.^72^. We extracted a vector shapefile of all islands for which we had any wild bee checklist information. In most cases we endeavored to associate each checklist with a single island entity. In cases where a single name from the bee database is associated with multiple land masses we extracted all the landmasses for subsequent calculations (e.g. Sao Tome & Principe, Saint Vincent & Grenadines, Wallis & Futuna, Antigua & Barbuda, Saint Pierre and Miquelon, Saint Kitts and Nevis, Solomon Islands). Duplicate island names and spatial mismatches were checked manually. Following this we harmonized the island names with the islands included in the Global Inventory of Floras and Traits (GIFT) database and extracted information on island area, isolation (distance to mainland), island category (oceanic, fragment or continental), archipelago and richness of angiosperms from GIFT (for more detail see^20^). Islands in the GIFT database were categorized by geological origin into oceanic (volcanic islands and atolls), continental shelf (land-bridge), and complex origin/continental fragments. Categorizing islands into three discrete types is a necessary simplification. Certain cases, such as the tectonic uplift of Cyprus or continental fragments like Rhodes and New Caledonia that underwent periods of total submergence, represent intermediate states where biological isolation was prioritized over strict geological origin. Specifically, a lack of historical land-bridge connections was used to classify islands as ‘oceanic’ in type Of the final 306 islands, there were 208 oceanic islands, 25 fragment islands and 73 continental islands (see Table S11 for a full breakdown by biome).

Species distributions and endemism data for 350,707 vascular plant species across 1,967 islands and 1,010 mainland regions were taken from Schrader et al.^9^ and then associated with islands for which we have bee species checklists. These data were originally sourced from the GIFT v.3.2 database using the GIFT R package^73^ and are taxonomically standardized according to the World Checklist of Vascular Plants (see^9,20^). In the few cases where we had bee checklist data for an island or archipelago that did not correspond to an island or archipelago in the GIFT database e.g. data aggregated at archipelago level in one but not the other, we calculated area and isolation directly from the aforementioned spatial map of islands^72^ and island specific literature was used to determine island category and archipelago.

We incorporated additional island-level environmental information by overlaying our island shapefile with a spatial global shapefile of ecoregions^33^ which is an update of the original WWF biogeographical regionalization of Olson et al.^74^. This dataset provides a globally standardized classification of terrestrial ecoregions, biomes and biogeographic realms, and was selected because it offers consistent, GIS-based borders that explicitly include oceanic islands, allowing reproducible assignment of islands based solely on spatial overlap. Although developed as a general terrestrial biodiversity classification rather than for specific taxa, the underlying climatic, floristic and evolutionary discontinuities that define these realms are expected to also be relevant for bees whose distributions are closely tied to vegetation structure and climatic regimes. Each island was assigned to a terrestrial biome and biogeographic realm. using the intersect function from the sf R package (v1.0-21^75^). Biome classifications were aggregated into broader categories (Boreal, Desert, Mediterranean, Temperate and Tropical) for comparative purposes (see Table S12) and the final selected biome was the dominant biome found on the island. Realm classifications were one of Afrotropic, Australasia, Indomalaya, Nearctic, Neotropic, Oceania, Palearctic. This allowed for consistent ecological context across islands and facilitated analyses of native bee richness by biome type and realm. To ensure comparability and ecological and biogeographical relevance, we limited our final dataset to islands with a minimum area of 5 km^2^ and located at least 1 km from the mainland. These thresholds helped remove islands with likely incomplete species inventories, high edge effects, and limited habitat heterogeneity, while ensuring a meaningful biogeographical break from the mainland.

### Database of all public bee records on islands and global bee checklist

To explore global patterns of bee sampling on islands we used the BeeBDC protocol outlined in Dorey et al.^70^ to extract all global records of bees on islands. Specifically, we downloaded all records of Anthophila from Global Biodiversity Information Facility^76^, Atlas of Living Australia^77^, and iDigBio. We corrected synonym names as above using the synonym lists from Dorey et al.^68^ and Orr et al.^4^. As part of the cleaning process the data was subset to only include names of species found in the island checklist dataset, ensuring incorrect records of valid species unlikely to occur on the island were removed. To spatially link bee occurrence records with islands, we reprojected all coordinates to match the island shapefile and performed a spatial join, retaining only records intersecting island polygons defined in the checklist dataset. To ensure consistency between occurrence and checklist data, all species listed in the island checklists were added as supplementary records to the public database, standardizing species richness values across sources. The final dataset comprised 663,189 georeferenced records, which were subsequently used to assess sampling effort and spatial coverage across islands included in the analysis. This dataset was used to calculate a proxy for sampling effort. We calculated sampling coverage across islands using the Chao & Jost^78^ method via the coverage function in the iNEXT3D R package (v1.08^79^). We used a species/island abundance (count of all occurrences per species per island) matrix of all records to estimate the proportion of the total individuals per island that belong to the observed species, using a non-parametric estimator that accounts for the frequency of rare species (singletons and doubletons). Because raw coverage can appear artificially high when based on very few records, we also applied a sample-size adjustment that scales coverage by the total number of occurrence records. This metric provides a standardized measure of sampling completeness across islands.

We used the Discover Life global checklist through the BeeBDC tool as our baseline source for global bee species richness. To generate an estimate of mainland-only species richness, we excluded all species known to occur on islands by comparing with the above checklists and by removing all island level records from the global checklist. The global checklist was also used to supplement endemism and island only species data by filtering out species only recorded on islands from the checklist and cross-referencing with the island dataset. The mainland checklist was also used to test island occurrence patterns to create a binary classification of all species as either occurring or not occurring on islands.

### Drivers of native bee richness

The island level checklists allow us to test the key drivers of species richness on islands. Here we provide a first analysis of the drivers of bee diversity on islands globally by focusing on the influence of island area and isolation^7^. We fit a linear mixed-effects model with log-transformed species richness as the response variable and log-transformed area, isolation and sampling effort, and island type and biome as predictors, using the lmer function from the lme4 R package (v1.1-37^80^). We log_10_-transformed native species richness after adding 1 to all values to accommodate islands with zero native bee species (N=7) while preserving them in the analysis. We began with a full model containing log_10_-transformed area, log_10_-transformed isolation (distance to nearest mainland), biome (categorical: five levels), island type (continental, fragment, oceanic), log_10_-transformed sampling effort (sample-size adjusted coverage), and all potential two- and three-way interactions among area, isolation, biome, and island type. Archipelago identity was included as a random intercept to account for spatial non-independence of islands within the same archipelago and to allow baseline richness to vary among archipelagos^81^. We used model selection via the dredge function from the MuMIn R package (v1.48.11^82^), with area and isolation retained as fixed effects in all candidate models, and ranked models by AIC_c_. We selected the best model based on the lowest AIC_c_; when more than one model was within ΔAIC_c_< 2, we selected the simplest model with the fewest degrees of freedom. The best-supported model (ΔAIC_c_> 2 to all other models) included:

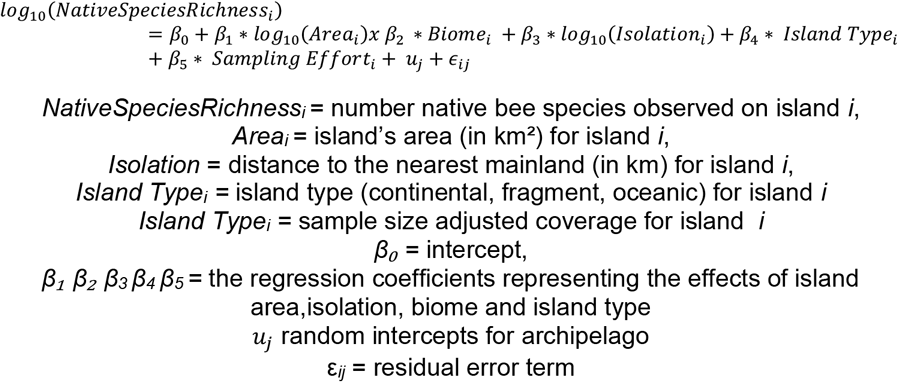

The final model included the interaction between area and biome, indicating that the effect of area on species richness varies among biomes. Island type and sampling effort were retained as a fixed effect. Two island groups for which coverage could not be calculated were dropped from the final model (Wallis and Futuna, Pitcairn Islands). We quantified variance explained by fixed effects alone using marginal R^2^ and variance explained by both fixed and random effects using conditional R^2,83^.To compare species–area relationship slopes among biomes, we extracted biome-specific slopes and their standard errors from the variance-covariance matrix of fixed effects. We conducted pairwise comparisons using z-tests, with p-values adjusted for multiple testing using the false discovery rate (FDR) method^84^. Biomes were assigned letter groupings to indicate homogeneous subsets (α = 0.05). Model predictions and confidence intervals were generated using the fixed-effects design matrix and visualized using ggplot2 (v3.5.2^85^). Finally, we fit the same best model for only oceanic and only continental and fragment islands, specifically to compare species-area relationship slopes among biomes of different island types.

### Drivers of bee endemism on islands and relationship with richness

To explore the drivers of endemic species richness and its relationship with total species richness across islands, we used a multi model approach mirroring our richness analysis. We fit a series of linear mixed-effects models using the lmer function from the lme4 R package (v1.1-37^79^). We analyzed patterns at the individual island scale using two measures of endemism as response variables: (1) single-island endemism (SIE), species restricted to individual islands, and (2) archipelago-level endemism (AE), species endemic to entire archipelagos. The archipelago-level approach captures archipelago-wide diversification processes while revealing which individual islands within archipelagos are endemism hotspots (see e.g.^45^).

We focused on islands supporting at least one single-island endemic (SIE ≥ 1; n = 101 islands). We started with the same full model as above for richness. In this subset of islands, all biomes are represented apart from boreal as they harboured no endemics. We used model selection via the dredge function from the MuMIn R package (v1.47.5^81^), with total species richness, area, and isolation retained as fixed effects in all candidate models (in Dataset 4, biome was additionally retained as a fixed effect), and ranked models by AIC_c_. We selected the best model based on the lowest AIC_c_; when more than one model was within ΔAIC_c_ < 2, we selected the simplest model with the fewest degrees of freedom. All models within ΔAIC_c_ < 2 are shown in Supplementary Table S7. The best-supported model included :

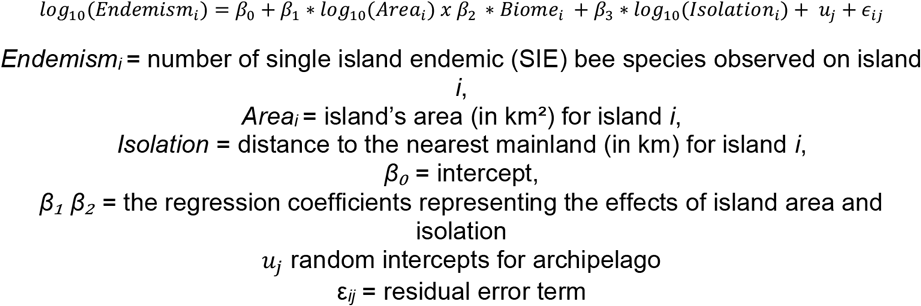

We additionally tested whether area-endemism slopes varied among biomes or were uniform. We explicitly tested this by comparing models with and without biome × area interactions using likelihood ratio tests.

To assess the generality of endemism patterns, we repeated analyses on three subsets: (1) oceanic islands only (n = 59), excluding potential pseudo-endemics resulting from recent land-bridge fragmentation; (2) endemic-rich islands (SIE > 5% of total richness; n = 81), focusing on systems where endemism is well-established. We also repeated all analyses for archipelago endemics (AE ≥ 1; n = 215) to confirm patterns were robust across endemism definitions.

Additionally, to formally test whether endemism patterns are spatially decoupled from richness patterns, we compared two nested models: (1) where endemism is predicted solely by richness: log_10_(SIE) ∼ log_10_(Richness) + (1|Archipelago); and (2) a where biome adds information beyond richness: log_10_(SIE) ∼ log_10_(Richness) + Biome + (1|Archipelago). If including biome significantly improves model fit (likelihood ratio test), this indicates that islands with similar richness have different endemism depending on biome. We then tested whether residual distributions differed among biomes using Kruskal-Wallis tests with post-hoc pairwise Wilcoxon rank-sum tests (Benjamini-Hochberg FDR correction).

### Bee-Plant Richness Mismatch Analysis

To assess whether bee diversity tracks angiosperm diversity across islands, we fitted a linear regression model with native bee richness as the response variable and native angiosperm richness as the predictor and tested for deviations of this relationship across biomes. We fit the model for the 181 islands for which we had both bee native richness data and reliable angiosperm richness data from the GIFT database^20^.

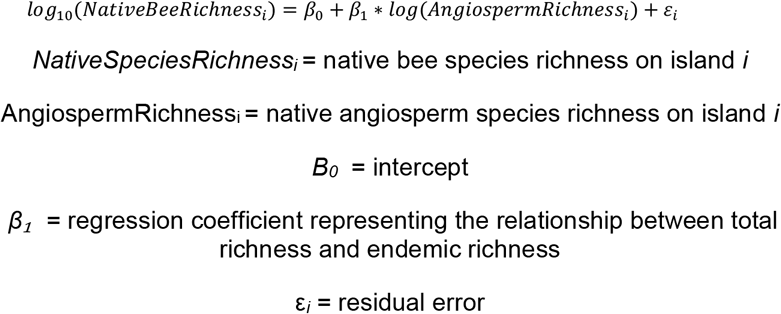

Residuals from this model represent the deviation of observed bee richness from values predicted by plant richness alone, with positive residuals indicating islands with more bee species than expected and negative residuals indicating fewer bee species than expected. We assessed whether residuals differed significantly among biomes using a Kruskal-Wallis test, as residuals were not normally distributed. We then conducted post-hoc pairwise comparisons using Wilcoxon rank-sum tests with Benjamini-Hochberg correction for multiple comparisons. To visualize groupings of biomes with similar bee-plant relationships, we generated letter groupings from the pairwise comparison p-values, biomes sharing a letter do not differ significantly (p ≥ 0.05). We visualized the distribution of residuals across biomes using combined violin and box plots, with median fold-change multipliers annotated on each biome.

## Acknowledgements

L.M. was supported by the Belgian F.R.S.-FNRS fellowships (Chargé de recherches) and the “BeeConnected” project (VI.Veni.222.141) which is financed by the Dutch Research Council (NWO). N.J.V. acknowledges the Belgian F.R.S-FNRS for financial support through the project “BeeGAPS: Bridging Data Gaps to Enhance Wild Bee Biodiversity Monitoring and Conservation in sub-Saharan Africa”(FNRS PDR T032125F). M.C.O. was supported by the Chinese Academy of Science’s President’s International Fellowship Initiative (2026PVC0122). H.K. acknowledges funding of research unit FOR2716 DynaCom (379417748) and Biodiversa+ BioMonI (533271599) from the German Research Foundation (DFG). Thank you to Thomas J. Wood for additional checklist data and discussions on the data and analysis. Also, thanks to Yilin Wu and Luis Valente for discussions about the analysis.

