## Supplemental Tables and Figures for "Global Patterns and Drivers of Bee Diversity and Endemism on Islands"

**Table S1 Introduced bees on islands globally.** Count shows the number of islands each species has been introduced on to.

| Species | Number of Islands |
|---|---|
| *Amegilla pulchra* | 4 |
| *Anthidium manicatum* | 4 |
| *Apis cerana* | 3 |
| *Apis mellifera* | 122 |
| *Bombus hortorum* | 3 |
| *Bombus hypnorum* | 1 |
| *Bombus jonellus* | 1 |
| *Bombus lucorum* | 1 |
| *Bombus pascuorum* | 1 |
| *Bombus pratorum* | 1 |
| *Bombus ruderatus* | 10 |
| *Bombus subterraneus* | 1 |
| *Bombus terrestris* | 7 |
| *Braunsapis puangensis* | 3 |
| *Ceratina arizonensis* | 3 |
| *Ceratina dentipes* | 11 |
| *Ceratina smaragdula* | 6 |
| *Euryglossina hypochroma* | 1 |
| *Euryglossina proctotrypoides* | 2 |
| *Hoplitis adunca* | 1 |
| *Hylaeus albonitens* | 6 |

| | |
|---|---|
| *Hylaeus asperithorax* | 3 |
| *Hylaeus perhumilis* | 2 |
| *Hylaeus strenuus* | 1 |
| *Hyleoides concinna* | 2 |
| *Lasioglossum helianthi* | 1 |
| *Lasioglossum impavidum* | 5 |
| *Lasioglossum microlepoides* | 1 |
| *Lithurgus scabrosus* | 16 |
| *Megachile chlorura* | 1 |
| *Megachile concinna* | 19 |
| *Megachile fullawayi* | 7 |
| *Megachile gentilis* | 4 |
| *Megachile lanata* | 22 |
| *Megachile otomita* | 1 |
| *Megachile rotundata* | 4 |
| *Megachile rufipennis* | 10 |
| *Megachile sculpturalis* | 1 |
| *Megachile umbripennis* | 15 |
| *Nomia melanderi* | 1 |
| *Osmia caerulescens* | 1 |
| *Xylocopa sonorina* | 12 |
| *Xylocopa violacea* | 1 |

**TableS2. Linear mixed-effects model of island bee species richness ($\log_{10}$-transformed).** Fixed effects include biome, island area, isolation, island type, and biome-specific area effects. Archipelago was included as a random intercept. Coefficients are shown with 95% confidence intervals. Biome terms are estimated relative to the reference biome. Biome × area terms represent deviations from the baseline species–area relationship. Significance levels: * $p < 0.05$, ** $p < 0.01$, *** $p < 0.001$.

| ***Term*** | ***Estimate*** | ***Std. Error*** | ***95% CI (lower)*** | ***95% CI (upper)*** |
|---|---|---|---|---|
| ***Intercept*** | 0.80 | 0.41 | -0.01 | 1.60 |
| ***mediterranean-type*** | -0.08 | 0.47 | -1.00 | 0.85 |

| | | | | |
|---|---|---|---|---|
| ***tropical*** | 0.02 | 0.41 | -0.78 | 0.82 |
| ***temperate*** | 0.34 | 0.42 | -0.49 | 1.17 |
| ***desert*** | 0.55 | 0.47 | -0.38 | 1.47 |
| ***Sampling effort*** | 0.22*** | 0.05 | 0.13 | 0.31 |
| ***Fragment Island*** | 0.07 | 0.10 | -0.12 | 0.27 |
| ***Oceanic island*** | -0.19** | 0.07 | -0.32 | -0.05 |
| ***Island area (log10)*** | 0.05 | 0.09 | -0.13 | 0.22 |
| ***Isolation (log10)*** | -0.13** | 0.04 | -0.22 | -0.05 |
| ***mediterranean-type:log_area*** | 0.45*** | 0.12 | 0.22 | 0.69 |
| ***tropical:log_area*** | 0.23* | 0.09 | 0.05 | 0.41 |
| ***temperate:log_area*** | 0.14 | 0.10 | -0.06 | 0.33 |
| ***desert:log_area*** | -0.03 | 0.13 | -0.30 | 0.23 |

**Table S3 Pairwise comparisons of species-area relationship slopes among biomes without accounting for sampling effort.** Biome-specific slopes (z) for the relationship between $\log_{10}$ island area ($km^2$) and $\log_{10}$ bee species richness were extracted from the best-supported mixed-effects model (isolation and sampling effort as additional fixed effects) and compared using z-tests. The slope difference represents the change in the species-area slope when moving from Biome1 to Biome2. Significant differences ($p < 0.05$) are indicated with an asterisk. P-values were adjusted for multiple testing using the false discovery rate (FDR) method. B = boreal, M = mediterranean-type, TC = tropical, TP = temperate, D = desert.

| Biome1 | Biome2 | Δ Slope | SE | z | p-value | FDR adj. p |
|---|---|---|---|---|---|---|
| **B** | **M** | -0.45 | 0.12 | -3.75 | 0.00*** | 0.00*** |

| | | | | | | |
|---|---|---|---|---|---|---|
| **B** | **TC** | -0.23 | 0.09 | -2.54 | 0.01* | 0.02* |
| **B** | **TP** | -0.14 | 0.10 | -1.38 | 0.17 | 0.19 |
| **B** | **D** | 0.03 | 0.13 | 0.24 | 0.81 | 0.81 |
| **M** | **TC** | 0.22 | 0.09 | 2.54 | 0.01* | 0.02* |
| **M** | **TP** | 0.32 | 0.09 | 3.48 | 0.00*** | 0.00** |
| **M** | **D** | 0.49 | 0.13 | 3.73 | 0.00*** | 0.00** |
| **TC** | **TP** | 0.10 | 0.05 | 1.89 | 0.06 | 0.09 |
| **TC** | **D** | 0.27 | 0.10 | 2.55 | 0.01* | 0.02* |
| **TP** | **D** | 0.17 | 0.11 | 1.54 | 0.12 | 0.16 |

**Table S4 Pairwise comparisons of species-area relationship slopes among biomes (a) oceanic island only and (b) continental and fragment islands only.** Biome-specific slopes (z) for the relationship between $\log_{10}$ island area and $\log_{10}$ bee species richness were extracted from the best-supported mixed-effects model and compared using z-tests. The slope difference represents the change in the species-area slope when moving from Biome1 to Biome2. Significant differences ($p < 0.05$) are indicated with an asterisk. P-values were adjusted for multiple testing using the false discovery rate (FDR) method. B = boreal, M = mediterranean-type, TC = tropical, TP = temperate, D = desert.

| ***(a) Oceanic islands*** | | | | | | |
|---|---|---|---|---|---|---|
| **Biome1** | **Biome2** | **Δ Slope** | **SE** | **z** | **p-value** | **FDR adj. p** |
| **B** | **M** | -0.30 | 0.26 | -1.14 | 0.25 | 0.36 |
| **B** | **TC** | 0.02 | 0.24 | 0.07 | 0.94 | 0.94 |
| **B** | **TP** | 0.17 | 0.25 | 0.66 | 0.51 | 0.56 |

| | | | | | | |
|---|---|---|---|---|---|---|
| **B** | **X** | 0.30 | 0.25 | 1.20 | 0.23 | 0.36 |
| **M** | **TC** | 0.31 | 0.12 | 2.57 | 0.01* | 0.03* |
| **M** | **TP** | 0.46 | 0.15 | 3.03 | 0.00** | 0.01* |
| **M** | **X** | 0.60 | 0.15 | 3.93 | 0.00*** | 0.00*** |
| **TC** | **TP** | 0.15 | 0.10 | 1.45 | 0.15 | 0.30 |
| **TC** | **X** | 0.29 | 0.10 | 2.78 | 0.01* | 0.02* |
| **TP** | **X** | 0.14 | 0.14 | 1.00 | 0.32 | 0.40 |

***(b) Continental islands***

| Biome1 | Biome2 | Δ Slope | SE | z | p-value | FDR adj. p |
|---|---|---|---|---|---|---|
| **B** | **M** | 0.50 | 1.30 | 0.39 | 0.70 | 0.87 |
| **B** | **TC** | 0.76 | 1.29 | 0.59 | 0.56 | 0.83 |
| **B** | **TP** | 0.81 | 1.29 | 0.63 | 0.53 | 0.83 |
| **B** | **X** | 0.73 | 1.33 | 0.55 | 0.58 | 0.83 |
| **M** | **TC** | 0.26 | 0.13 | 1.94 | 0.05 | 0.26 |
| **M** | **TP** | 0.31 | 0.14 | 2.23 | 0.03 | 0.26 |
| **M** | **X** | 0.23 | 0.33 | 0.68 | 0.50 | 0.83 |
| **TC** | **TP** | 0.05 | 0.08 | 0.58 | 0.56 | 0.83 |
| **TC** | **X** | -0.03 | 0.31 | -0.10 | 0.92 | 0.92 |
| **TP** | **X** | -0.08 | 0.32 | -0.25 | 0.80 | 0.89 |

**Table S5 Pairwise comparisons of species-area relationship slopes among biomes without accounting for sampling effort.** Biome-specific slopes (z) for the relationship between $\log_{10}$ island area ($km^2$) and $\log_{10}$ bee species richness were extracted from the best-supported mixed-effects model and compared using z-tests. The slope difference represents the change in the species-area slope when moving from Biome1 to Biome2. Significant differences ($p < 0.05$) are indicated with an asterisk. P-values were adjusted for multiple testing using the false discovery rate (FDR) method. B = boreal, M = mediterranean-type, TC = tropical, TP = temperate, D = desert.

| Biome1 | Biome2 | Δ Slope | SE | z | p-value | FDR adj. p |
|---|---|---|---|---|---|---|
| B | M | -0.46 | 0.13 | -3.69 | 0.00*** | 0.00** |
| B | TC | -0.27 | 0.10 | -2.83 | 0.00** | 0.01* |
| B | TP | -0.15 | 0.10 | -1.44 | 0.15 | 0.19 |
| B | D | 0.00 | 0.14 | 0.02 | 0.99 | 0.99 |
| M | TC | 0.20 | 0.09 | 2.17 | 0.03* | 0.04* |
| M | TP | 0.32 | 0.10 | 3.33 | 0.00*** | 0.00** |
| M | D | 0.47 | 0.14 | 3.45 | 0.00*** | 0.00** |
| TC | TP | 0.12 | 0.05 | 2.28 | 0.02* | 0.04* |
| TC | D | 0.27 | 0.11 | 2.52 | 0.01* | 0.02* |
| TP | D | 0.15 | 0.11 | 1.33 | 0.19 | 0.21 |

**Table S6. Drivers of single-island endemism (SIE) and archipelago endemism (AE).** Linear mixed-effects models relating bee endemism to island characteristics. Values are standardized regression coefficients (β) with 95% confidence intervals. Archipelago was included as a random intercept. in all models. Significance: *** $p < 0.001$; ** $p < 0.01$; * $p < 0.05$; · $p < 0.1$. Given that many islands harbor zero endemic species, we fit models across four nested datasets to understand how drivers of endemism vary depending on island characteristics and total native species richness. (i) All islands (including islands with zero endemics); (ii) islands with endemics (SIE ≥ 1 or AE ≥ 1); (iii) oceanic islands only (with endemics) and (iv) endemic-rich islands

(>5% endemic species). M = mediterranean-type, TC = tropical, TP = temperate, D = desert. Boreal islands harboured no endemic species and were excluded from this analysis.

| Model | Term | Estimate (ß) | SE | 95% CI |
|---|---|---|---|---|
| **SIE >= 1**<br>n=101<br>Marginal $R^2$ = 0.50<br>Conditional $R^2$ = 0.50 | Intercept | -0.47* | 0.24 | [-0.939, -0.002] |
| | Island area (log10) | 0.35*** | 0.04 | [0.278, 0.42] |
| | Isolation (log10) | -0.06 | 0.07 | [-0.201, 0.074] |
| **Oceanic Only (SIE >= 1)**<br>n=59<br>Marginal $R^2$ = 0.28<br>Conditional $R^2$ = 0.28 | Intercept | -0.45 | 0.27 | [-0.989, 0.087] |
| | Island area (log10) | 0.25*** | 0.05 | [0.143, 0.356] |
| | Isolation (log10) | 0.03 | 0.09 | [-0.145, 0.207] |
| **Endemic-rich (SIE> 5%)**<br>n=81<br>Marginal $R^2$ = 0.65<br>Conditional $R^2$ = 0.65 | Intercept | -0.25 | 0.25 | [-0.745, 0.242] |
| | Island area (log10) | 0.39*** | 0.03 | [0.318, 0.451] |
| | Isolation (log10) | -0.15* | 0.07 | [-0.294, -0.002] |
| **AE >=1**<br>n=215<br>Marginal $R^2$ = 0.55 | Intercept | -0.08 | 0.33 | [-0.739, 0.578] |
| | Biome: TC | -0.25 | 0.34 | [-0.916, 0.413] |

| | | | | |
|---|---|---|---|---|
| Conditional $R^2$ = 0.73 | Biome: TP | -0.18 | 0.39 | [-0.951, 0.596] |
| | Biome: D | 0.41 | 0.41 | [-0.386, 1.215] |
| | Sampling effort (log10) | 0.17** | 0.05 | [0.064, 0.273] |
| | Island area (log10) | 0.22* | 0.10 | [0.025, 0.412] |
| | Isolation (log10) | -0.02 | 0.06 | [-0.149, 0.102] |
| | Biome TC × Area | 0.16 | 0.10 | [-0.039, 0.363] |
| | Biome TP × Area | 0.09 | 0.12 | [-0.135, 0.318] |
| | Biome D × Area | -0.17 | 0.14 | [-0.454, 0.111] |

**Table S7. Alternative candidate models (ΔAICc < 2) for single-island endemism (SIE) and archipelago endemism (AE) across island datasets.** Models shown here represent all supported alternatives to the best-supported models presented in Table S6, and ΔAICc values are calculated relative to those best models. Effect sizes correspond to standardized fixed-effect coefficients; blank cells indicate that a term was not included in that model. Archipelago was included as a random intercept in all models. Model sets are shown for: (i) SIE on islands with ≥1 endemic species, (ii) SIE on endemic-rich islands (>5% endemic species), and (iii) AE on islands with ≥1 endemic species. No alternative models were supported for oceanic islands (ΔAICc < 2). **+** indicates included in model and **-** indicates not included in the model.

| Predictor | SIE >=1 a | SIE >=1 b | SIE >=1 c | Endemic-rich (SIE> 5%) | AE >=1 |
|---|---|---|---|---|---|

| | | | | | |
|---|---|---|---|---|---|
| **(Intercept)** | -0.47 | -0.43 | -0.45 | -0.25 | 0.13 |
| **Biome** | + | + | - | 0 | + |
| **Island area (log10)** | 0.34 | 0.32 | 0.34 | 0.39 | 0.16 |
| **Isolation (log10)** | -0.14 | -0.12 | -0.05 | -0.15 | 0.00 |
| **Sampling Effort** | - | 0.13 | 0.06 | - | 0.17 |
| **Island Classification** | - | - | - | - | + |
| **Biome × Area** | - | | - | - | + |
| **Biome x Isolation** | - | - | - | - | - |
| **df** | 8 | 9 | 6 | 5 | 14 |
| **logLik** | -53.55 | -52.50 | -56.28 | -34.11 | -82.56 |
| **AICc** | 124.7 | 125 | 125.5 | 79 | 195.2 |
| **delta** | 0.97 | 1.26 | 1.75 | 0.75 | 1.23 |
| **weight** | 0.17 | 0.15 | 0.117 | 0.35 | 0.23 |

**Table S8. Fixed effects and model comparison of native richness ~ endemism models for single island endemism (SIE) and archipelago endemism (AE).** (A) Model output from SIE and AE models. Significance: *** $p < 0.001$, ** $p < 0.01$, * $p < 0.05$. with and without biome. (B) ANOVA model comparison between sIE and AE models with and without Biome included as a fixed effect. $\Delta R^2$ represents change in marginal $R^2$; $\chi^2$ values from likelihood ratio tests (ML estimation). M = mediterranean-type, TC = tropical, TP = temperate, D = desert. M is the baseline biome. Boreal islands harboured no endemic species and were excluded from this analysis.

| ***(a) Model output*** |
|---|

| Response | Model | term | Estimate (ß) | SE | 95% CI |
|---|---|---|---|---|---|
| SIE | SIE ~Richness | Intercept | -0.311* | 0.12 | [-0.549, -0.074] |
| | SIE ~Richness | Final richness (log10) | 0.635*** | 0.07 | [0.491, 0.779] |
| | SIE ~Richness + Biome | Intercept | -1.257*** | 0.18 | [-1.623, -0.891] |
| | SIE ~Richness + Biome | Final richness (log10) | 0.787*** | 0.07 | [0.65, 0.924] |
| | SIE ~Richness + Biome | Biome: TC | 0.877*** | 0.14 | [0.596, 1.158] |
| | SIE ~Richness + Biome | Biome: TP | 0.677*** | 0.18 | [0.31, 1.044] |
| | SIE ~Richness + Biome | Biome: X | 0.913*** | 0.23 | [0.45, 1.375] |
| AE | AE ~Richness | Intercept | -0.415*** | 0.07 | [-0.561, -0.269] |
| | AE ~Richness | Final richness (log10) | 0.862*** | 0.03 | [0.807, 0.916] |
| | AE ~Richness + Biome | Intercept | -1.177*** | 0.13 | [-1.426, -0.929] |

| | | | | | |
|---|---|---|---|---|---|
| | AE ~Richness + Biome | Final richness (log10) | 0.876*** | 0.03 | [0.82, 0.932] |
| | AE ~Richness + Biome | Biome: TC | 0.92*** | 0.13 | [0.664, 1.177] |
| | AE ~Richness + Biome | Biome: TP | 0.888*** | 0.14 | [0.613, 1.164] |
| | AE ~Richness + Biome | Biome: X | 0.951*** | 0.15 | [0.658, 1.245] |

**b) ANOVA model comparison of above models**

| Response | N | AIC | AIC w/Biome | Delta AIC | R2 | R2 w/Biome | Delta R2 | $\chi^2$ | df | P value |
|---|---|---|---|---|---|---|---|---|---|---|
| **SIE** | 101 | 140.65 | 120.40 | 20.24 | 0.40 | 0.58 | 0.18 | 33.09 | 3 | <0.001 |
| **AE** | 215 | 70.07 | 45.86 | 24.21 | 0.54 | 0.66 | 0.12 | 40.04 | 3 | <0.001 |

**Table S9 Pairwise comparisons of species-area relationship slopes among biomes for single island archipelago endemism (AE).** Biome-specific slopes (z) for the relationship between $\log_{10}$ island area ($km^2$) and $\log_{10}$ AE were extracted from the best-supported mixed-effects model (isolation was the additional fixed effect) and compared using z-tests. The slope difference represents the change in the species-area slope when moving from Biome1 to Biome2. Significant differences ($p < 0.05$) are indicated with an asterisk. P-values were adjusted for multiple testing using the false discovery rate (FDR) method. M = mediterranean-type, TC = tropical, TP = temperate, D = desert. Boreal islands harboured no endemic species and were excluded from this analysis.

| Biome1 | Biome2 | Δ Slope | SE | z | p-value | FDR adj. p |
|---|---|---|---|---|---|---|
| **M** | **TC** | -0.16 | 0.10 | -1.56 | 0.11 | 0.22 |

| | | | | | | |
|---|---|---|---|---|---|---|
| **M** | **TP** | -0.09 | 0.12 | -0.79 | 0.42 | 0.42 |
| **M** | **D** | 0.17 | 0.14 | 1.37 | 0.23 | 0.34 |
| **TC** | **TP** | 0.07 | 0.07 | 1.02 | 0.29 | 0.35 |
| **TC** | **D** | 0.33 | 0.11 | 3.28 | 0.00** | 0.01* |
| **TP** | **D** | 0.26 | 0.12 | 2.38 | 0.03 | 0.09 |

**Table S10. P-values from pairwise comparisons of biome residuals from angiosperm ~ native bee richness model, using Wilcoxon rank sum exact test.** P-value adjustment method following the Benjamini-Hochberg procedure.

| | Boreal | Desert | Tropical | Temperate |
|---|---|---|---|---|
| **Desert** | 0.46 | - | - | - |
| **Tropical** | 0.11 | 0.11 | - | - |
| **Temperate** | 0.03* | <0.001*** | | - |
| **Mediterranean-type** | <0.001*** | <0.001*** | <0.001*** | <0.001*** |

**Table S11 Breakdown of island category by biome.**

| | continental | fragment | oceanic |
|---|---|---|---|
| **Boreal** | 2 | 0 | 5 |
| **Mediterranean-type** | 10 | 7 | 27 |
| **Tropical** | 41 | 9 | 136 |
| **Temperate** | 19 | 7 | 19 |
| **Desert** | 1 | 2 | 21 |

**Table S12 Original biome classifications from Dinerstein et al. (2017) and simplified aggregations used for analysing island area/isolation relationships.**

| Original Biome | Biome |
|---|---|
| Boreal Forests/Taiga | Boreal |
| Tundra | Boreal |
| Deserts & Xeric Shrublands | Desert |
| Mediterranean Forests, Woodlands & Scrub | Mediterranean-type |
| Temperate Broadleaf & Mixed Forests | Temperate |
| Temperate Conifer Forests | Temperate |
| Temperate Grasslands, Savannas & Shrublands | Temperate |
| Tropical & Subtropical Coniferous Forests | Tropical |
| Tropical & Subtropical Dry Broadleaf Forests | Tropical |
| Tropical & Subtropical Grasslands, Savannas & Shrublands | Tropical |
| Tropical & Subtropical Moist Broadleaf Forests | Tropical |
| Mangroves | Tropical |

### Figures

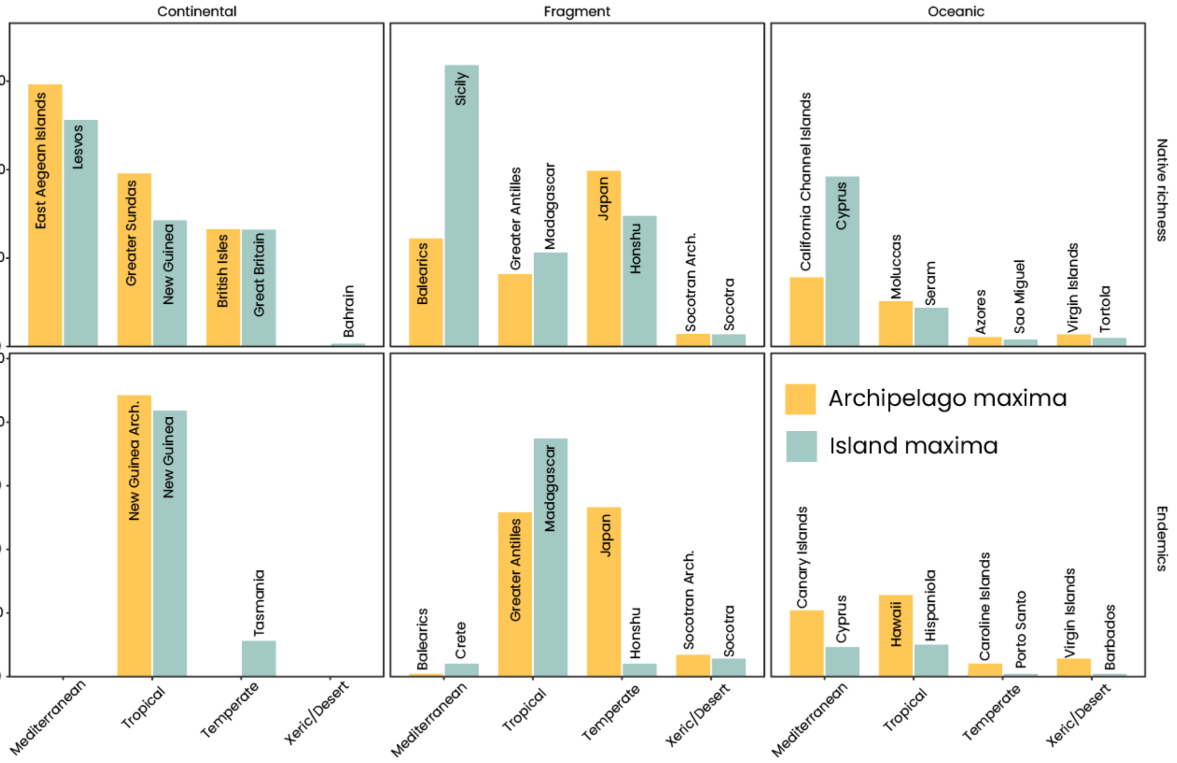


**Figure S1. Island- and archipelago-level biodiversity extremes in native richness across Biomes and Island Types.** Bars show the highest island-level and highest archipelago-level values for three biodiversity metrics, native species richness, single-island endemics (SIE), and archipelago endemics (AE), calculated separately for each Biome and island type combination. Island maxima identifies the single island within each Biome–TypeModel group with the highest recorded value, while archipelago maxima identifies the archipelago whose dominant biome and island type (determined by total island area) exhibits the greatest total richness or endemism. Labels above bars indicate the name of the corresponding island or archipelago. Values for each metric are shown in separate rows, and island types are shown in separate columns. This figure highlights how peak biodiversity contributions differ between individual islands and their broader archipelagic context across global biomes.

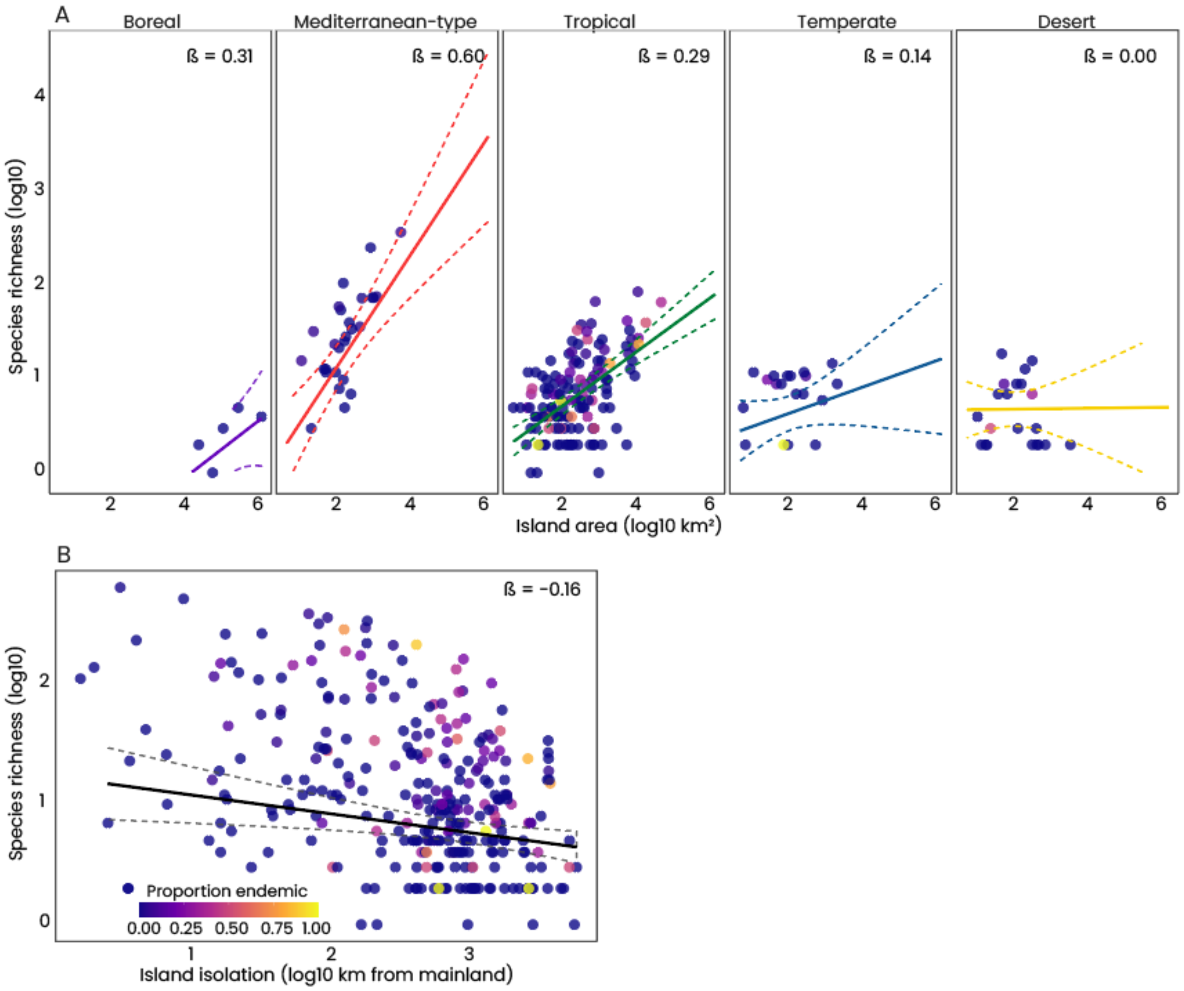


**Figure S2. Drivers of island bee species richness for oceanic islands.** (A) Biome-specific species–area relationships. Solid lines represent fitted slopes from mixed-effects models, with dashed lines indicating 95% confidence intervals. Slopes (β) are reported for each biome. Point colour indicates the proportion of endemic species per island. Labelled islands are the top 10% most species-rich islands within each biome (values in parentheses show total native bee richness). (B) Relationship between island isolation (log$_{10}$ distance to mainland) and native bee species richness. Point colour indicates the proportion of endemic species per island.

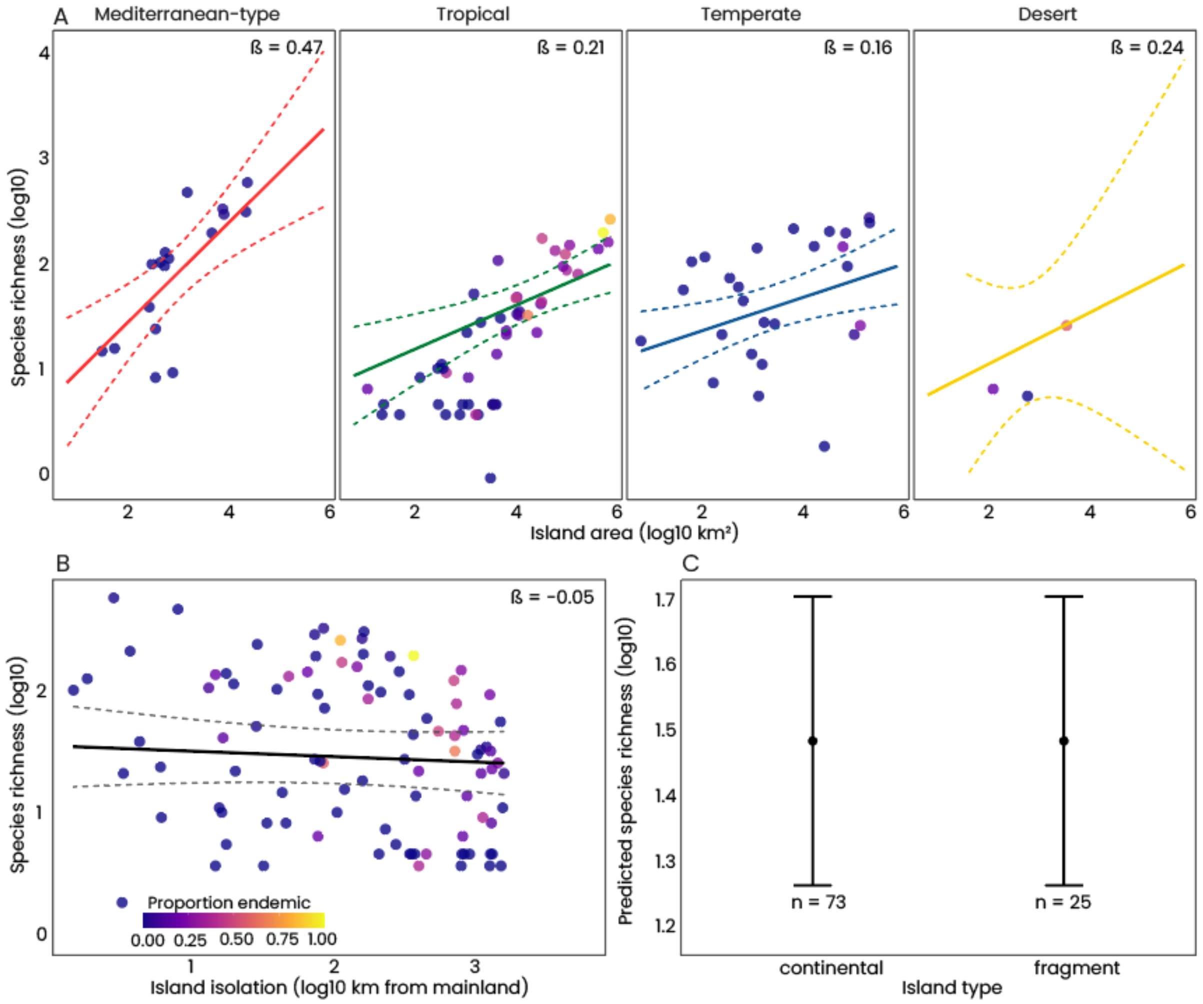


**Figure S3. Drivers of island bee species richness for continental and fragment islands.** (A) Biome-specific species–area relationships. Solid lines represent fitted slopes from mixed-effects models, with dashed lines indicating 95% confidence intervals. Slopes (β) are reported for each biome. Point colour indicates the proportion of endemic species per island. Labelled islands are the top 10% most species-rich islands within each biome (values in parentheses show total native bee richness). (B) Relationship between island isolation ($log_{10}$ distance to mainland) and native bee species richness. Point colour indicates the proportion of endemic species per island. (C) Predicted bee species richness by island type (continental, fragment, oceanic) from the full mixed-effects model, holding other predictors constant. Circles indicate the model-predicted means ± 95% confidence intervals, with the number of islands shown below each category.

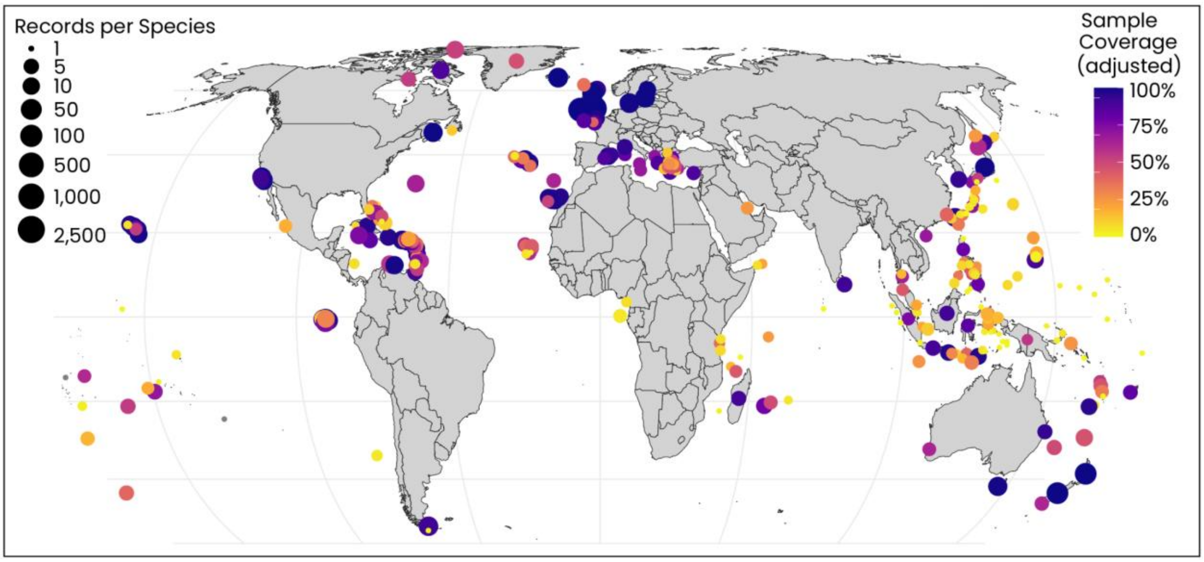


**Figure S4. Global sampling coverage and effort for island bee research.** Map shows the geographic distribution of bee sampling effort and completeness across islands worldwide. Point color indicates sample coverage (completeness estimate, ranging from 0% in yellow to 100% in purple), reflecting the proportion of estimated true diversity captured by sampling. Circle size represents the number of individual records (specimens or observations see methods section "Database of all public bee records on islands and global bee checklist" ) collected per species on each island, with larger circles indicating greater sampling effort. Islands with dark purple coloring and large point sizes represent well-sampled regions with high confidence in richness estimates. Yellow and orange points represent under-sampled islands where true diversity is likely underestimated. Coverage was calculated s across islands using the Chao & Jost[77] method via the coverage function in the iNEXT3D R package (v1.08[78]). We used a species/island abundance (count of all occurrences per species per island) matrix of all records to estimate the proportion of the total individuals per island that belong to the observed species, using a non-parametric estimator that accounts for the frequency of rare species (singletons and doubletons). Because raw coverage can appear artificially high when based on very few records, we also applied a sample-size adjustment that scales coverage by the total number of occurrence records. This metric provides a standardized measure of sampling completeness across islands. This metric provides a standardized measure of sampling completeness across islands.

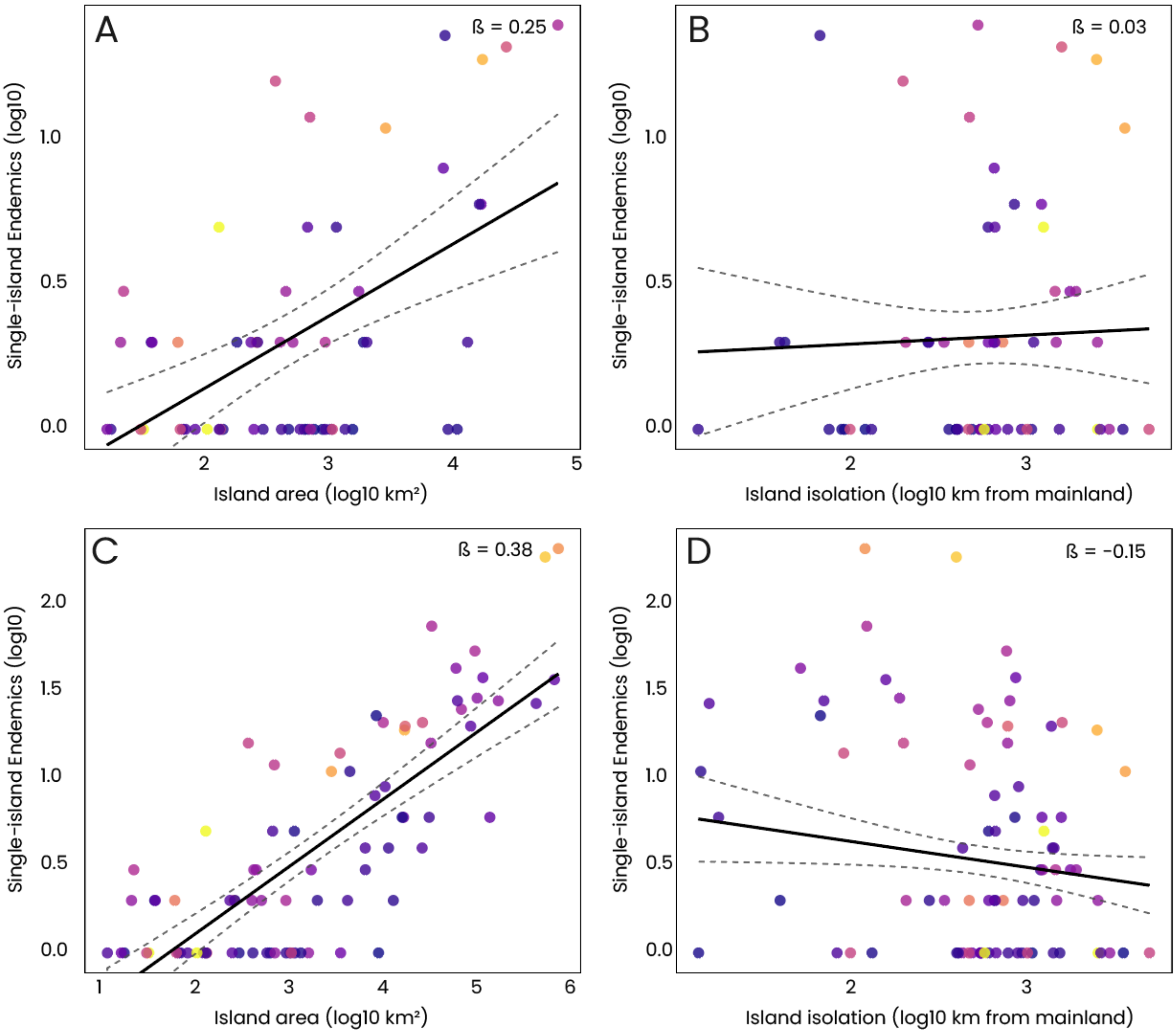


**Figure S5. Drivers of single-island endemic bee richness in oceanic and endemism rich islands.** (A) Relationship between island area ($\log_{10}$ km²) and single-island endemic (SIE) species richness for oceanic islands with at least one single-island endemic (n = 59). Black line shows the overall trend with 95% confidence interval (gray shading). Circles are colored by the proportion of native species that are endemic. (B) Relationship between island isolation ($\log_{10}$ distance to nearest mainland, km) and SIE richness for oceanic islands. (C) Relationship between island area ($\log_{10}$ km²) and single-island endemic (SIE) species richness for islands where endemics comprise more than 5% of native bee richness (n = 81). Black line shows the overall trend with 95% confidence interval (gray shading). Circles are colored by the proportion of native species that are endemic. (D) Relationship between island isolation ($\log_{10}$ distance to nearest mainland, km) and SIE richness for endemic-rich islands. Black line shows the overall trend with 95% confidence interval. Circle colors as in panel A.

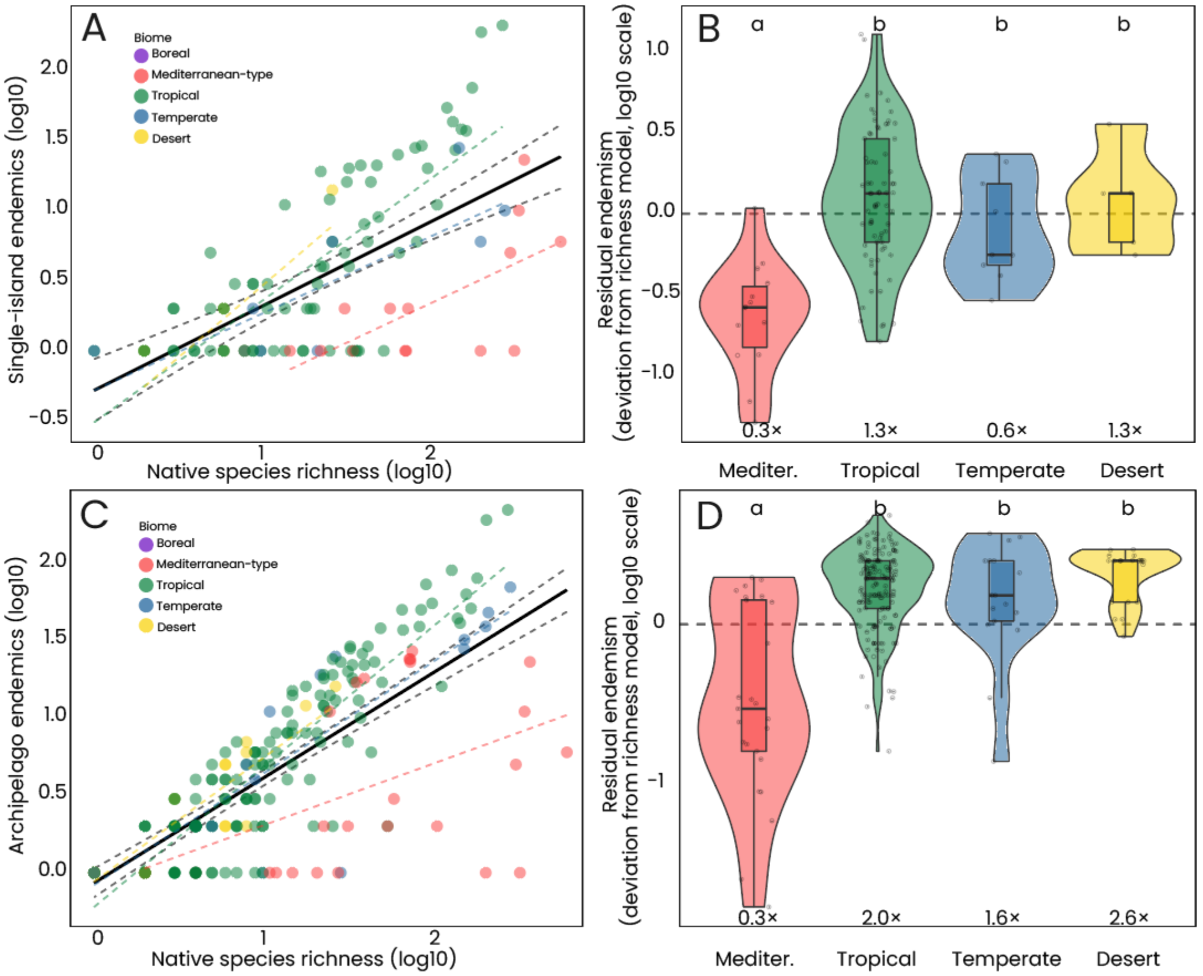


**Figure S6. Relationship between island bee endemism and native species richness across biomes.** (A) Relationship between total native species richness ($log_{10}$) and single-island endemic (SIE) richness ($log_{10}$), colored by biome. The dashed black line shows the overall linear trend (±95% CI, gray shading); dashed colored lines show biome-specific relationships. Islands with at least one SIE (n = 101). Panel A is the same as Fig. 3d. (B) Residuals from a linear mixed model of $log_{10}$(SIE) ~ $log_{10}$(native richness) with archipelago as a random effect, shown by biome. Values above zero indicate more SIEs than predicted by richness alone; values below zero indicate fewer. Numbers show the median fold-change relative to model predictions. Biomes differed significantly in residual SIE richness (Kruskal–Wallis test; letters denote groups with significantly different residuals based on pairwise Wilcoxon tests with Benjamini–Hochberg correction, $p < 0.05$). (C) As in A but for archipelago endemics (AE; islands with at least one AE, n = 101). (D) As in B but for residuals from the AE richness model.
